# mNeonGreen-tagged fusion proteins and nanobodies reveal localization of tropomyosin to patches, cables, and contractile actomyosin rings in live yeast cells

**DOI:** 10.1101/2022.05.19.492673

**Authors:** Tomoyuki Hatano, Tzer Chyn Lim, Ingrid Billault-Chaumartin, Anubhav Dhar, Ying Gu, Teresa Massam-Wu, Sushmitha Adishesha, Luke Springall, Lavanya Sivashanmugam, William Scott, Masanori Mishima, Sophie G Martin, Snezhana Oliferenko, Saravanan Palani, Mohan K. Balasubramanian

**Author notes:** TH and TCL contributed equally to this work. IBC, AD, YG, and TMW contributed equally to this work and are listed alphabetically. Correspondence: Mohan Balasubramanian.

## Abstract

Tropomyosins are structurally conserved α-helical coiled-coil dimeric proteins that bind along the length of filamentous actin (F-actin) in fungi and animals. Tropomyosins play essential roles in the stability of actin filaments in non-muscle cells and are essential for calcium regulation of myosin II contractility in the muscle. Despite the crucial role of tropomyosin in actin cytoskeletal regulation, *in vivo* investigations of tropomyosin are limited, mainly due to the suboptimal live cell imaging tools currently available in many organisms. Here, we report mNeon-Green (mNG) tagged tropomyosin, with native promoter and linker length configuration, that clearly reports tropomyosin localization and dynamics in *Schizosaccharomyces pombe* (Cdc8), *Schizosaccharomyces japonicus* (Cdc8), and *Saccharomyces cerevisiae* (Tpm1 and Tpm2), *in vivo* and in isolated *S. pombe* cell division apparatuses. We extended this approach to also visualize the mammalian TPM2 isoform. Finally, we generated a camelid-nanobody against *S. pombe* Cdc8, which mimics the localization of mNG-Cdc8 in vivo without significantly influencing cell growth and dynamics of actin cytoskeleton. Using these tools, we report the presence of tropomyosin in previously unappreciated patch-like structures in fission and budding yeasts, show flow of tropomyosin (F-actin) cables to the cytokinetic actomyosin ring, and identify rearrangements of the actin cytoskeleton during mating. These powerful tools and strategies will aid better analyses of tropomyosin and actin cables *in vivo*.

## Introduction

Tropomyosins are coiled-coil α-helical dimeric proteins that bind and strengthen actin filaments. These proteins are 160-280 amino acids in length (in ~ integer multiples of 40) in which approximately 40 amino acids from tropomyosin spans an actin monomer along the actin filament (Gimona, 2008; Gunning et al., 2015; Holmes and Lehman, 2008). Tropomyosins assemble into unidirectional head-to-tail polymers along an actin filament and this polymerization increases affinity for actin filaments ~ 100-fold (Gimona, 2008; Gunning et al., 2015; Holmes and Lehman, 2008). They are widely present in fungal and animal lineages, but clear tropomyosin homologs have not been identified in other phyla / kingdoms (Gunning et al., 1997; Gunning et al., 2015; Lin et al., 1997; Martin and Gunning, 2008; Perry, 2001; Pruyne, 2008). In the muscle, tropomyosins regulate actin-myosin II interaction in response to Ca^2+^ release (Brown and Cohen, 2005; Chalovich et al., 1981; Gergely, 1974; Perry, 2001; Spudich and Watt, 1971; Szent-Gyorgyi, 1975; Wakabayashi, 2015). They assemble along with filamentous actin and troponins into “thin filaments” whose interaction with the myosin II motors located in “thick filaments” generates contractile forces. In animal non-muscle cells, tropomyosins play crucial roles in cytoskeletal organization, cell polarity, and cytokinesis and are detected in stress fibers, lamellipodia, cleavage furrow, and the cell cortex (Gunning et al., 2015; Lin et al., 1997). Through work largely in budding and fission yeasts, fungal tropomyosins have been shown to localize to long formin-generated actin cables, the cytokinetic apparatus, and the fusion focus, an actin-rich zone observed during mating and fusion of cells of opposite mating types (Alioto et al., 2016; Balasubramanian et al., 1992; Drees et al., 1995; Dudin et al., 2017; Gunning et al., 2015; Huckaba et al., 2004; Liu and Bretscher, 1992; Pruyne, 2008; Skau et al., 2011; Skau and Kovar, 2010; Wloka et al., 2013; Yang and Pon, 2002). Imaging of fixed wild-type cells with antibodies shows that the *S. pombe* tropomyosin also localizes to actin patches (Balasubramanian et al., 1992; Skoumpla et al., 2007). However, none of the available fluorescent protein tools for live imaging detect tropomyosin in patches.

Despite the essentiality of tropomyosins for cell viability and for stabilization of formin-generated actin filaments, investigation of its dynamic properties in live cells are limited. It is largely due to the sub-optimal fluorescent probes available for *in vivo* imaging of tropomyosins, especially in the fungi. In this work, we have designed and generated a new mNeon-Green (Shaner et al., 2013) (mNG)-tropomyosin probe that detects actin cables, actomyosin rings, and fusion foci in one or more of three different yeasts (*S. pombe, S. japonicus*, and *S. cerevisiae*). We also extended this approach to investigate tropomyosin dynamics in mammalian cells, using human RPE cells as an example. Finally, we developed camelid nanobodies against *S. pombe* tropomyosin encoded by the *cdc8* gene, which expands the toolkit to study tropomyosin function in this yeast and provides a general strategy for the study of *in vivo* dynamics of tropomyosin in other organisms.

## Results

### Live imaging of mNeon-Green (mNG) tagged *S. pombe* Cdc8-tropomyosin

In *Schizosaccharomyces pombe*, the *cdc8* gene encodes an essential 161 amino acid tropomyosin (Balasubramanian et al., 1992). This protein is essential for actin cable stability, cell fusion focus formation, and cytokinetic actomyosin ring (CAR) assembly, and thus, for cell mating and cytokinesis (Billault-Chaumartin and Martin, 2019; Christensen et al., 2019; Dudin et al., 2017; Pelham and Chang, 2001; Skau et al., 2009). Consistently, staining of wild-type *S. pombe* cells (Balasubramanian et al., 1992; Palani et al., 2019; Skoumpla et al., 2007) with antibodies raised against Cdc8 reveals its localization in cables that run along the long axis of the cell and cytokinetic actomyosin rings (Figure 1A). Cdc8 is also detected in patches at the cell ends, which in several cases are connected to the actin cables (Skoumpla et al., 2007) (Figure 1A). Cdc8 has been visualized in live cells as a fusion with a green fluorescent protein (GFP), under the control of an attenuated thiamine-repressible *nmt1* promoter (Billault-Chaumartin and Martin, 2019; Johnson et al., 2014; Maundrell, 1993). GFP-Cdc8 is clearly detected in the CAR, weakly in actin cable, and not detected in patches (Pelham and Chang, 2002) (Skoumpla et al., 2007) and therefore hasn’t been used to investigate patch or cable dynamics.

**Figure 1:**
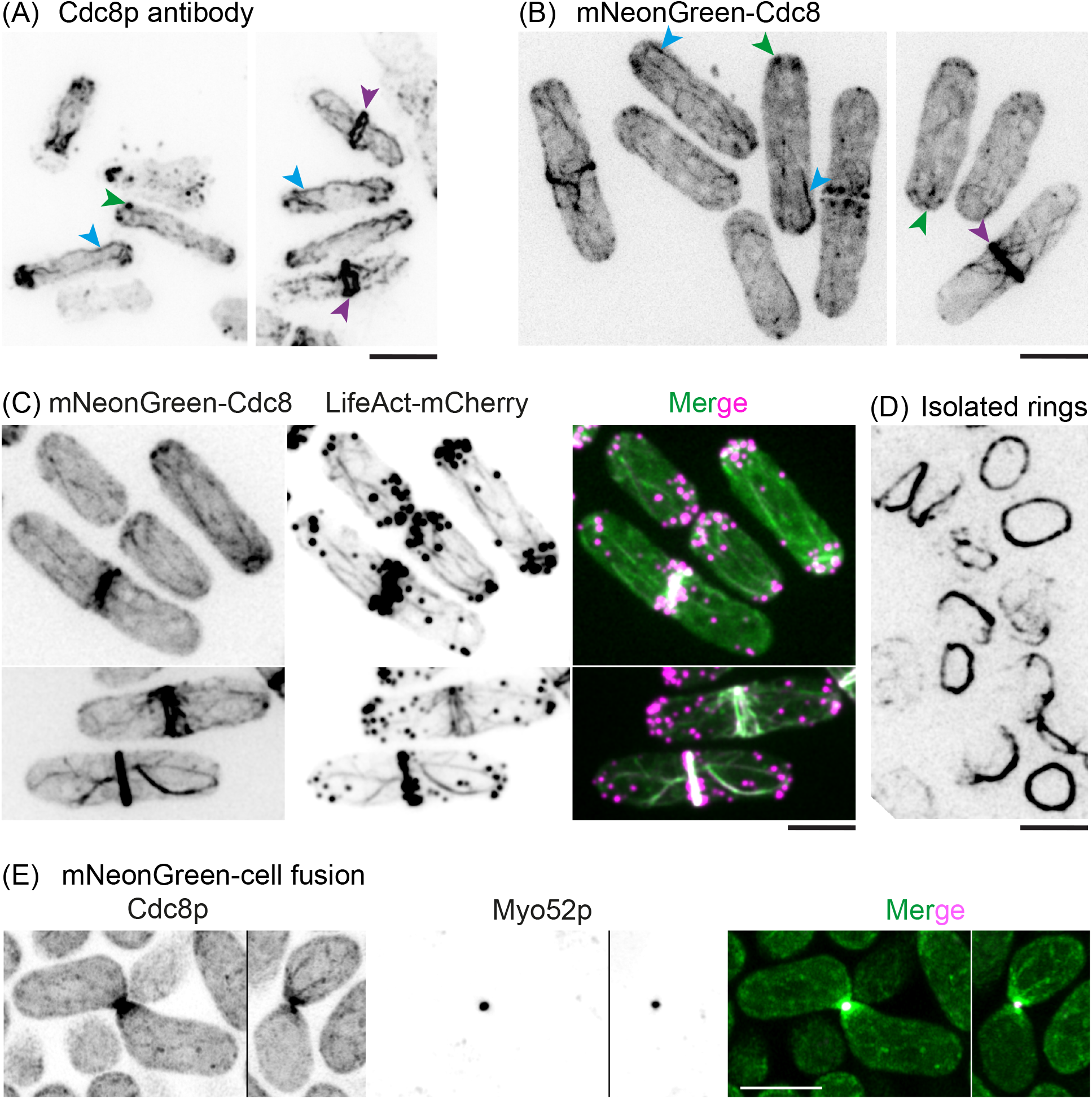
Cdc8-tropomyosin localization to patches, cables, the CAR and the fusion focus in *S. pombe*. (A) Wild-type cells were fixed and treated with antibodies against Cdc8 to visualize Cdc8-Tropomyosin. (B) Cdc8 was N-terminally tagged with mNeonGreen to visualize its localization in patches, cables and cytokinetic ring. Green arrowheads: mNG-Cdc8 patches. Blue arrowheads: mNG-Cdc8 cables and purple arrowheads: mNG-Cdc8 in the CAR. (C) Cells coexpressing mNG-cdc8 (left) and LifeAct-mCherry (mid) shows the co-localization of actin and Cdc8. Merged fluorescence images shows that while actin cables and Cdc8 cables visibly colocalize, only a subset of F-actin patches contained Cdc8 signal. (D) Cytokinetic rings were isolated from cells expressing mNG-Cdc8 and the image shows stable association of mNG-Cdc8 in CARs within cell ghosts. (E) Airyscan2 images of mating cells expressing mNG-Cdc8 (green) and Myo52-tdTomato (magenta), which labels the fusion focus. mNG-Cdc8 strongly accumulates at the fusion focus and labels actin cables, which can be seen emanating from the fusion focus and at other positions in the cells. Scale bars are 5μm.

To facilitate *in vivo* study of tropomyosin dynamics, we therefore set out to make an improved probe. Three parameters (*cdc8* promoter driving fusion gene expression, an mNG fluorescent protein, and a 40 amino acid flexible linker (Hussain et al., 2018) between mNG and Cdc8) helped generate a fission yeast strain in which we were able to visualize Cdc8 in all three structures (Figure S1A). Through live imaging of this strain (Figure 1B), as with antibody staining of fixed cells, mNG-Cdc8 was detected in small patches, actin cables (some of which were connected to actin patches), and the actomyosin ring (which was connected to a meshwork of actin cables). Although expression of mNG-Cdc8 on the top of native Cdc8 did not cause any overt dominant phenotype (Figure S1B), as with the previous GFP-Cdc8 probe, it was not fully functional and was unable to sustain life when present as the sole copy. Nevertheless, this experiment established that Cdc8 is a component of actin patches, cables, and the CAR.

To investigate whether mNG-Cdc8 colocalized with F-actin, we imaged a strain expressing mNG-Cdc8 and, LifeAct-mCherry as a marker for F-actin (Figure 1C). As expected, F-actin was detected in patches, cables, and CARs and mNG-Cdc8 colocalised with actin cables and the CAR. Interestingly, although present in multiple small patches, mNG-Cdc8 was present in fewer patches compared with F-actin. Previous work has shown that Cdc8-tropomyosin competes with the actin filament cross-linker fimbrin for actin binding (Christensen et al., 2017). Consistently, in cells lacking fimbrin (*fim1*Δ), mNG-Cdc8 was more obvious in actin patches (Figure S1C).

Next, we tested if mNG-Cdc8 was retained in CARs isolated from *S. pombe* cells, which should facilitate the investigation of tropomyosin (and F-actin) dynamics during CAR constriction *in vitro*.To this end, mNG-Cdc8 expressing cells were spheroplasted and permeabilized with detergent to isolate cell ghosts carrying the CAR. mNG-Cdc8 was stably associated with the CAR held within permeabilized cell ghosts (Mishra et al., 2013) (Figure 1D). This experiment established that mNG-Cdc8 was not only potentially useful to investigate tropomyosin dynamics in live cells, but also in isolated CARs.

In *S. pombe*, upon nitrogen starvation, cells of opposite mating types polarize and grow towards each other culminating in the formation of the fusion focus, a formin-assembled structure underlying the concentration of secretory vesicles transported by the myosin V Myo52. This is followed by cell-cell fusion, nuclear fusion, meiosis, and sporulation. In mating cells (Figure 1E), mNG-Cdc8 was prominently detected in the fusion focus, where it decorated a larger region than that marked by Myo52. It also decorated longer cables that appeared to emanate from the zone of cell-cell contact, as well as fine speckles.

The mNG-Cdc8 strain generated in this work provided a far superior signal to noise compared to the currently available GFP-Cdc8 strains, in which the *GFP-cdc8* fusion gene is expressed under control of the thiamine-repressible *nmt*41/42 promoter directly fused to the Cdc8-tropomyosin coding sequence without an intervening linker (Pelham and Chang, 2002) (Figure S2). Collectively, these experiments established that mNG-Cdc8 is a reliable marker that can be used to investigate Cdc8-tropomyosin dynamics in patches, cables, CAR, and during cell fusion.

### mNG-Cdc8 is suitable for long-duration time-lapse imaging in vegetative and mating *S. pombe* cells

Having generated the mNG-Cdc8 strain, we tested its usefulness in several time-lapse imaging experiments. Wild-type cells expressing mNG-Cdc8 were imaged every 3 seconds for ~ 30 minutes using a spinning disk confocal microscope. mNG-Cdc8 did not undergo significant photobleaching over this time and therefore mNG-Cdc8 enabled us to see tropomyosin dynamics in *S.pombe* at higher time resolution.

In interphase cells, multiple small patches of mNG-Cdc8 were detected, some of which showed directional movement from the cell ends towards the cell middle along mNG-Cdc8 cables (Figure 2A and Supplemental Movie 1). In mitotic cells, mNG-Cdc8 localized as short filaments at the division site presumably loading on to Formin-Cdc12 induced actin filaments assembled from cytokinetic nodes (Vavylonis et al., 2008) (Figure 2B, top panel, and Supplemental Movie 2). Furthermore, non-medially located mNG-Cdc8 cables were transported to the cell middle where they incorporated into the forming CAR as shown previously for F-actin cables detected using Lifeact-GFP (Huang et al., 2012) Figure 2B, bottom panels and Supplemental movie 2). To demonstrate the Cdc8 flow toward the cell middle, a kymograph was generated by a line segmentation of the cells along the long axis of rod-shaped cell in maximum intensity projected z-stack images (Figure 2C). Transport of non-medially located mNG-Cdc8 cables into the CAR was also observed in the highly elongated *cdc25-22* cells (Figure S3 and Supplemental Movie 3 and 4) in which these movements were more obvious. In cells undergoing septation, mNG-Cdc8 cables and mNG-Cdc8 aggregates were found to leave the CAR concomitant with its constriction (Figure 2D and Supplemental movie 5). We were also able to observe mNG-Cdc8 in isolated CARs (Figure 2E), which upon addition of ATP underwent constriction (Supplemental movies 6). This is indeed consistent with the earlier study showing that tropomyosin is associated with actin bundles that are expelled during CAR constriction (Huang et al., 2016).

**Figure 2:**
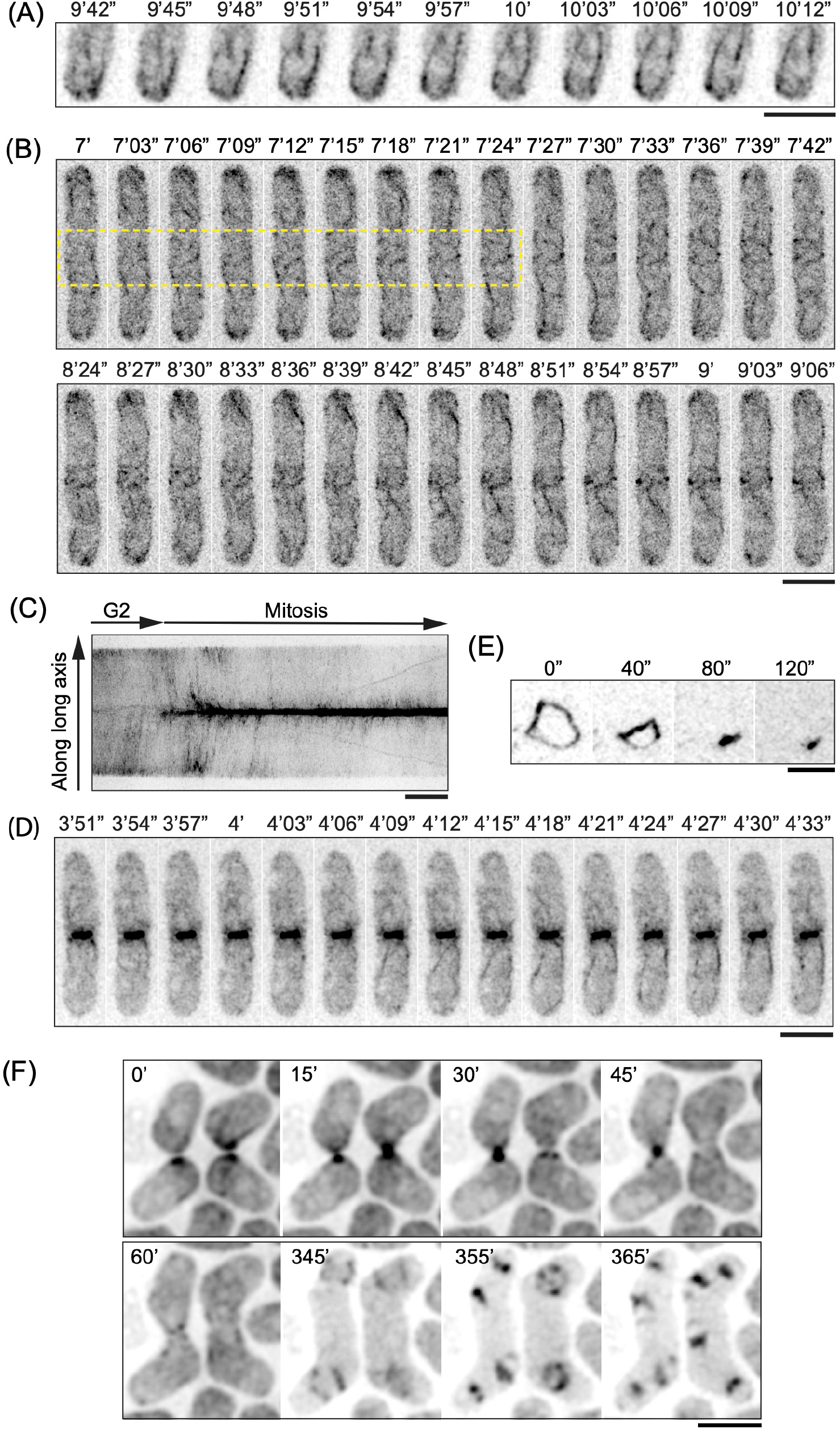
Visualization of mNG-Cdc8 by time-lapse imaging. (A) Time-lapse images of wildtype cells expressing mNG-Cdc8 at interphase. Cdc8 exists as patches and cables originate from cell tips. Patches can be seen moving on the cables as the cables elongate. (B) Time-lapse images of CAR assembly in wild-type cells expressing mNG-Cdc8. The top panel shows events in a 42 second window in which mNG-Cdc8 assembles into short filaments at the division site and in long cables elsewhere (within the yellow box). The bottom panel shows movement of non-medial actin cables into the forming CAR. (C) Kymograph of a line parallel to the long axis of a cell at interphase entering mitosis – mNG-Cdc8 signals are found at cell ends, subsequently move towards mid-cell and accumulate and form the CAR. (D) Time-lapse images showing mNG-Cdc8 behaviour in a CAR within a cell ghost undergoing constriction following ATP addition. The CAR within the cell ghost constricted in 120 seconds. (E) Time-lapse images of wild-type cells during ring constriction in which mNG-Cdc8 cables are seen to be expelled from the CAR. (F) Time-lapse epifluorescence images of mating cells expressing mNG-Cdc8. Two pairs of fusing cells are shown, where mNG-Cdc8 decorates the fusion focus (15-45 min), as well as some actin cables before fusion (0 min). The fusion focus disassembles upon fusion (60 min). After the diploid zygote has undergone meiosis, mNG-Cdc8 accumulates on the meiotic actin rings that form during sporulation (345-365 min). Scale bars are 5μm.

Next, we tested if mNG-Cdc8 could be used to investigate tropomyosin dynamics in cells undergoing mating and sporulation. We used epifluorescence imaging to acquire images of mNG-Cdc8 every 5 min for 12 h (Figure 2F and supplemental movie 7). mNG-Cdc8 localized at growth projections, strongly accumulating at the fusion site, until cell-cell fusion when the signal disappeared, as previously described for the fusion focus (Dudin et al., 2015). A strong mNG-Cdc8 signal re-appeared several hours later, after the diploid zygote had undergone meiosis, with mNG-Cdc8 accumulating on the meiotic actin rings that form during sporulation (Yan and Balasubramanian, 2012).

Collectively, this work established that mNG-Cdc8 is a powerful reporter for time-lapse imaging of Cdc8-tropomyosin dynamics in vegetative and sexually reproducing *S. pombe* cells.

### mNG-Cdc8 and mNG-Tpm1/2 as tools to visualize tropomyosin in *S. japonicus, S. cerevisiae*, and human RPE cells

Following the successful visualization of *S. pombe* Cdc8-tropomyosin using the mNG fusion, we investigated if the same strategy (Promoter-mNG-40 amino acid linker-Cdc8 or Tpm1/2) led to visualization of tropomyosin containing cellular structures in *S. japonicus* and *S. cerevisiae*.Previous work in *S. cerevisiae* has reported Tpm1 and Tpm2 in the CAR and actin cables in cells fixed and stained with antibodies (Pruyne et al., 1998), but they haven’t been visualized in cables in live cells. The 161-amino acid single tropomyosin has not been visualized in live *S. japonicus* cells.

First, we constructed a *S. japonicus* strain (Figure S4A) in which the coding sequences for mNG-Cdc8 [P_cdc8_-mNG-40aa linker-cdc8] were integrated into the *ura4* locus as a second copy. As in *S. pombe*, this strain did not show any dominant deleterious growth or division defects and colony formation rate was comparable to that of wild-type cells (Figure S4B).

As in *S. pombe, S. japonicus* mNG-Cdc8 localized to speckles / patches, cables, the CAR, and at the fusion focus during mating (Figure 3A). In time lapse imaging, mNG-SjCdc8 was detected in non-medially placed cables which were transported and incorporated into the CAR (Figure 3B and Supplemental movie 8). Furthermore, mNG-Cdc8 cables were expelled from the constricting CAR as seen with F-actin during CAR constriction in *S. japonicus* (Huang et al., 2016) (Figure 3B and Supplemental movie 8). The incorporation of non-medial mNG-Cdc8 cables into the CAR and its expulsion during CAR constriction were even better resolved in the elongated *cdc25^ts^* mutants after G2 arrest and release (Figure S5 and Supplemental movie 9). As in *S. pombe*, the mNG-SjCdc8 fluorescence did not bleach rapidly and therefore mNG-SjCdc8 is a useful tool to investigate tropomyosin dynamics in *S. japonicus*.

**Figure 3:**
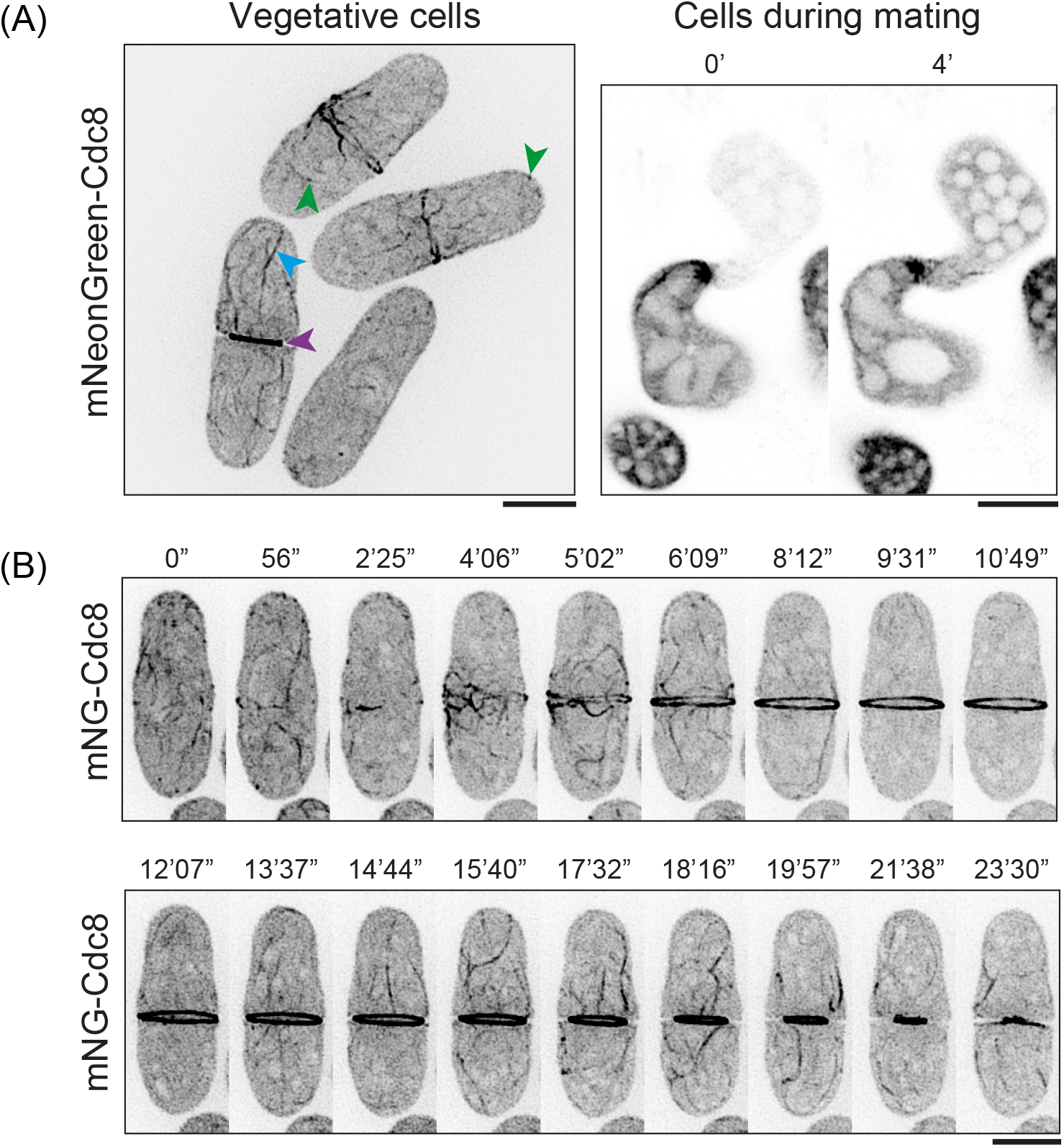
Visualization of mNG-Cdc8 in *S. japonicus*. (A) Left: *S. japonicus* cells with mNG-Cdc8 patches (green arrowhead), cables (blue arrowhead) and CAR (purple arrowhead). Right: Cdc8 patches and cables are seen in *S. japonicus* cells during the mating process. (B) Time-lapse images of CAR assembly and constriction in a *S. japonicus* cell. Note that mNG-Cdc8 assembles at the division site. Longer cables of mNG-Cdc8 also incorporate into the CAR. During constriction, mNG-Cdc8 in cables are expelled from the CAR. Scale bars are 5μm.

Next, we constructed mNG tagged tropomyosin in budding yeast *S. cerevisiae*, in which the actin cytoskeleton has been extensively characterized. There are two tropomyosin encoding genes in *S. cerevisiae*, Tpm1 (199 amino acids) (Liu and Bretscher, 1989) and Tpm2 (161 amino acids) (Drees et al., 1995; Wang and Bretscher, 1997). Tpm1 abrogation causes loss of detectable actin cables in the cell which leads to morphological defects. Furthermore, cells lacking Tpm1 and Tpm2 are inviable (Drees et al., 1995), establishing that as in *S. pombe*, tropomyosin function is essential for S. cerevisiae viability. We made the appropriate strains to image Tpm1 and Tpm2, which expressed the appropriate fusion proteins: P_tpm1_-mNG-40 aa linker-Tpm1 and P_Tpm2_-mNG-40 aa linker-mNG-Tpm2 (Figure S6A). Cells expressing mNG-Tpm1 and mNG-Tpm2 grew and formed colonies almost identically to wild-type cells (Figure S6B) and did not show any overt morphological or cell division defects.

mNG-Tpm1 was detected in patches and in long cables starting near the bud neck and migrating into the mother cell (Figure 4Ai and B and Supplemental movie 10). During cytokinesis, Tpm1 was detected at the division site in the CAR as reported previously (Okada et al., 2021) (Figure 4Ai and C and Supplemental movie 11). mNG-Tpm2 was detected as a strong focus at the bud site and in cables fainter than those visualized with mNG-Tpm1 (Figure 4Aii and D and Supplemental movie 12). However, it was clearly detected in the CAR (Figure 4Aii and E and Supplemental movie 13). Interestingly, as in *S. pombe*, Tpm1 and Tpm2 were detected in multiple patches in cells deleted for *SAC6*, the gene encoding budding yeast fimbrin (Figure S6C). Tpm1/Tpm2 patches were detected in both the mother and daughter cells. This work establishes that mNG-Tpm1 is a reliable tool to visualize Tpm1 in cables and the CAR, and mNG-Tpm2 appears better in the detection of Tpm2 in the CAR, while only weakly detecting Tpm2 in cables.

**Figure 4:**
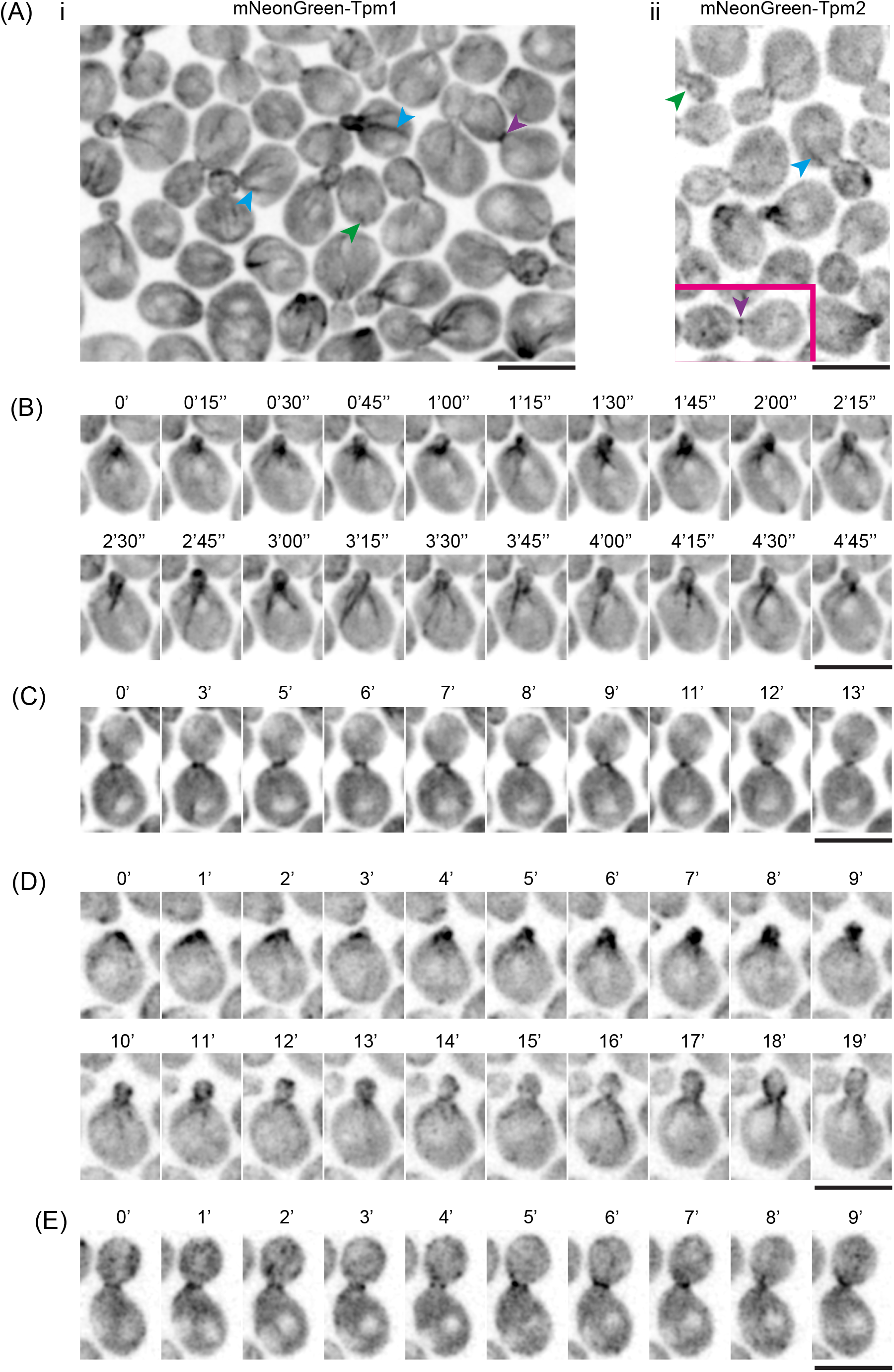
Visualization of mNG-Tpm1 and mNG-Tpm2 in *S. cerevisiae*. (A) Images of a field of cells expressing (i) mNG-Tpm1 and (ii) mNG-Tpm2. Patches are indicated with green arrowheads, cables with blue arrowheads and actomyosin rings with purple arrowheads. (B) Time-lapse (min) images revealing dynamic cables of mNG-Tpm1 in cables (C). Time-lapse images of mNG-Tpm1 dynamics duting cytokinesis Scale bars are 5μm. (D) Time-lapse (min) images revealing dynamic cables of mNG-Tpm2 in cables (E). Time-lapse images of mNG-Tpm2 dynamics during cytokinesis Scale bars are 5μm.

Finally, we tested if the mNG-40 aa linker – tropomyosin-based visualization of tropomyosin worked in mammalian cells. To this end, we generated pcDNA3.1 carrying mNG-40 aa linker – TPM2 (a highly expressed non-muscle tropomyosin isoform splice variant expressing a 284 amino acid protein), in which expression was driven from the cytomegalovirus (CMV) promoter, and the expressed fusion gene was as follows P_CMV_ – mNG – 40 aa linker – HsTPM2. When transiently transfected into human RPE cells, we were able to detect clear contractile stress fibers, which colocalized with F-actin in fixed cells co-stained with Rhodamine-conjugated phalloidin, consistent with past work (Appaduray et al., 2016; Tojkander et al., 2011; Yang et al., 2020).

These experiments in 4 organisms / cell types established that mNG-40 aa linker – tropomyosin is a reliable marker to investigate tropomyosin (and associated actin filament) dynamics.

### A camelid nanobody faithfully reports *S. pombe* Cdc8p-tropomyosin localization

In recent years single domain camelid nanobodies have become powerful tools to investigate protein localization and function (Kirchhofer et al., 2010). These nanobodies are relatively small (~ 15kDa), soluble, and suitable for various light and super-resolution microscopic approaches, including to directly visualize protein conformational states (Keller et al., 2019; Kirchhofer et al., 2010; Liu et al., 2020). We sought to make nanobodies against *S. pombe* Cdc8 as a proof of concept to test efficacy of nanobodies in investigating tropomyosin dynamics. To this end, we purified recombinant dimeric Cdc8-tropomyosin, which was used to commercially select interacting nanobodies through phage display screening followed by yeast two-hybrid screening (Moutel et al., 2016). We obtained a panel of seven nanobodies, which were screened through additional yeast two-hybrid screening (Figure S8A) and their ability to detect Cdc8-tropomyosin *in vivo* as N- or C-terminal fusions expressed from one of three promoters (ADH11, ADH21, and ADH81). The entire panel of nanobodies and their full characterization will be reported elsewhere. One of the nanobodies (Nanobody 5 – Nb5) was efficient in detecting Cdc8 *in vivo*. AlphaFold2 (Jumper et al., 2021; Jumper and Hassabis, 2022) structural predictions revealed that this nanobody (Nb5) potentially binds both chains of the dimeric Cdc8 via its complementarity determining regions (CDR), which is predicted to form hydrogen bonds with E89, E92, E94, T97, and R103 of Cdc8 and interact with a hydrophobic patch between the two chains of Cdc8 (L90, L91, and L95) (Figure 5A). Note AlphaFold2 prediction of interaction between Cdc8-tropomyosin dimer compared with a nonspecific Nb against β-catenin (Braun et al., 2016) as a control (Figure 5A and supplemental movie 14). When Nb5 was expressed as a C-terminal mNG fusion, it bound all structures detected previously using Cdc8 antibodies and mNG-Cdc8 fusion (Figure 5B). In interphase cells, Nb5 detected patches and cables that followed the cell long axis and in mitotic cells it detected the CAR. In time-lapse studies, Nb5 detected flow of Cdc8 (actin) cables from non-medial locations into the CAR (Supplemental movie 15). Finally, as in the case of mNG-Cdc8, Nb5 also decorated Cdc8-tropomyosin in the fusion focus and the signal from mNG-Cdc8 and Nb5-mNG were comparable (Figure 5D). We note that in mating assays, expression of Nb5 led to mild fusion defects and cell lysis. Collectively, this work provides evidence that the camelid nanobody technology can be successfully applied to the investigation of tropomyosin dynamics.

**Figure 5:**
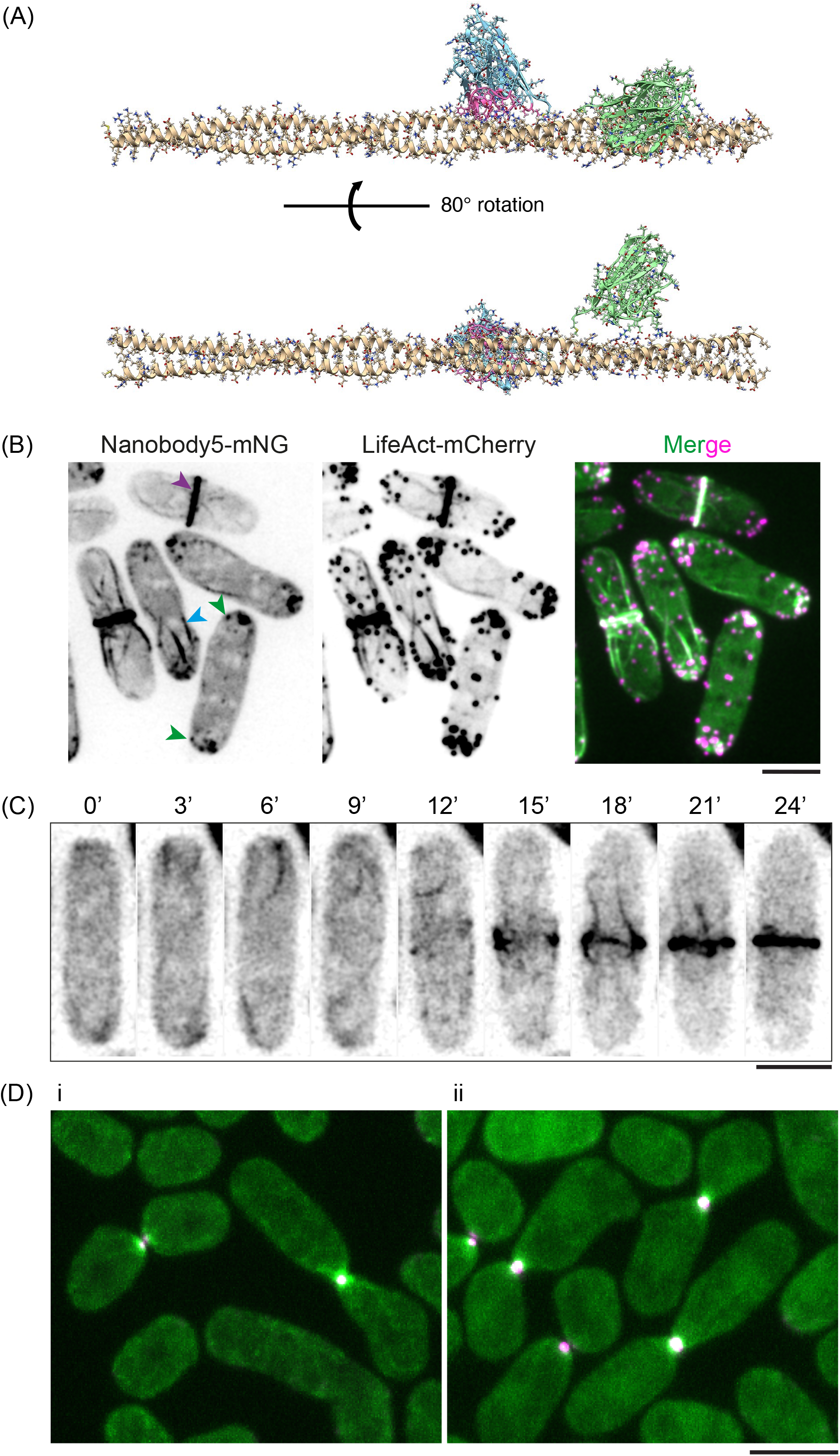
Visualization of Cdc8 with targeting-nanobody in *S. pombe*. (A-B) Alphafold2 prediction of binding between dimeric *S. pombe* Cdc8-tropomyosin with Nb5. Nb5 (blue) is predicted to bind, via its Complementarity Determining Regions (CDRs; shown in pink), amino acids 86-103 on Cdc8 (golden color dimer). Alphafold2 prediction of a non-specific nanobody targeting a peptide from β-catenin (PMID: 26791954) (green) is also shown. (C) *S. pombe* cells expressing Nanobody 5 (Nb5) fused to mNG (Nb5-mNG) detect Cdc8-tropomyosin in patches (green arrowheads), cables: blue arrowheads, and the CAR (purple arrowheads). (D) Time-lapse images of dynamics of Nb5-mNG as a marker for Cdc8-tropomyosin during CAR assembly. (E) Spinning disk microscopy images of *S. pombe* mating cells. Panel i – Mating cells expressing mNG-Cdc8 (green) and Myo52-tdTomato (magenta). The merge is shown in white. Panel ii – Mating cells expressing Nb5-mNG (green) and Myo52-tdTomato (magenta). The merge is shown in white. Scale bars are 5μm.

## Discussion

In this work we have developed tools to visualize tropomyosins in 4 different organisms / cell types using an mNG-fusion strategy. These yeast strains and plasmids for tropomyosin visualization will be made available to the community. Given that the mNG-40 aa linker – tropomyosin works in 4 scenarios, we believe this strategy will be transferrable to investigate tropomyosin in other fungi and metazoans. Although *S. pombe* cells expressing mNG-Cdc8 as their sole copy of Cdc8 were inviable (data not shown), this is not uncommon in that several essential cytoskeletal proteins do not support viability when expressed as the sole tagged copy (e.g. actin, tubulin, ESCRTIII), and are typically expressed as tagged second copies (Chen et al., 2012; Snaith et al., 2010). Nevertheless, study of fusions of these cytoskeletal proteins with various fluorescent proteins have enriched our understanding of the cytoskeleton. We note that although, mammalian TPM2 was expressed transiently in our work, the successful tagging opens the possibility of generating stably expressing cell lines and transgenic gene replacement animals expressing the mNG – 40 aa linker-TPM fusion gene under native promoter sequences. Future work should evaluate how the mNG-TPM2 fusion we describe compares with those described elsewhere in side-by-side experiments (Appaduray et al., 2016; Tojkander et al., 2011; Yang et al., 2020).

We also describe a workflow, and proof-of-concept in *S. pombe*, to develop Cdc8-tropomyosin nanobodies, an mNG tagged version of which we use to visualize tropomyosin *in vivo*, thereby expanding the tool kit to image tropomyosin. In addition to directly developing nanobodies against tropomyosin from other organisms as a primary strategy, an attractive additional possibility is to “evolve” the *S. pombe* tropomyosin nanobodies (based on similarity to the binding epitope) to identify Nb5-variants that recognize tropomyosin from other organisms. We have determined that Nb5 potentially binds amino acids 86-103 on Cdc8-tropomyosin (the molecular characterization of this interaction and those of the other 6 nanobodies with Cdc8-tropomyosin will be reported elsewhere).

### What new insights have we gained and what more can be addressed using these new tools?

First, although fungal tropomyosins have been studied for their roles in actin cables and the CAR, we have shown using our mNG-40 aa linker-tropomyosin fusions, that they localize to patches in three wild-type yeast species. A previous study demonstrated Cdc8-tropomyosin localization in patches at the cell ends in live fimbrin-defective mutants (Christensen et al., 2017), although Cdc8-tropomyosin decorated patches have been observed in wild-type cells only using immunofluorescence microscopy on fixed and permeabilized cells. The fact that Cdc8-tropomyosin was detected in patches in live wild-type *S. pombe* cells using two different probes established that Cdc8-tropomyosin, during normal physiology, may only be partly out-competed by fimbrin and may have a function in patches. This localization mirrors the weak, occasional localization of the formin For3 at actin patches, which is mostly outcompeted from patches by capping proteins (Billault-Chaumartin and Martin, 2019). Cdc8-Tropomyosin in patches may thus represent the imperfect sorting of actin-binding proteins. However, the presence of Cdc8-Tropomyosin only in a subset of actin patches suggests that it may alter or regulate actin patch function, perhaps playing important temporal roles in patch biogenesis and function. The localization of mNG-Tpm1/mNG-Cdc8 to patches in *S. cerevisiae* and *S. japonicus* raises the possibility that tropomyosin may have functional roles in actin patches in other fungi as well. It is also possible that formins use severed actin filaments from actin patches to elongate linear actin cables and the patches may represent sites from which tropomyosins are loaded onto actin filaments. The dynamics and function of tropomyosin in actin patches and how it relates to other actin patch components and endocytosis and how it relates to formins and actin cable elongation can now be studied using the tools we have generated.

Second, previous work on CAR assembly has used non-physiological probes to visualize CAR Factin assembly (such as CHD, Utrophin, or LifeAct), which influences kinetics of F-actin polymerization and depolymerization (Huang et al., 2012; Vavylonis et al., 2008). Therefore, it has been debatable if the F-actin non-medial flow toward the ring was physiological. mNG-Cdc8 is observed in *de novo* generated filaments at the division site and in non-medial cables that are transported to the CAR in *S. pombe*. These signals are lost upon treatment with the actin polymerization inhibitor latrunculin A (Ayscough et al., 1997), establishing that the mNG-Cdc8 can be a valuable tool to investigate actin cables. Our observation with Cdc8-tropomyosin dynamics provides direct evidence for two mechanisms in operation (direct F-actin nucleation at the cell division site from cytokinetic nodes and transport of non-medial F-actin cables) during CAR F-actin assembly. The rate of assembly, the extent of turnover, and the velocity of cable movement can now be investigated. The ability to use the superior mNG-Cdc8 in CARs in cell ghosts also allows the investigation of Cdc8-tropomyosin and actin filament dynamics during myosin-II dependent CAR constriction (triggered by ATP addition to isolated CARs).

Third, mNG-Cdc8 provides superior imaging of the actin cytoskeleton during mating and meiosis. While Cdc8 had previously been described at the fusion focus (Dudin et al., 2017; Kurahashi et al., 2002), mNG-Cdc8 reveals a more extensive signal that extends over a larger zone than that occupied by the type V myosin Myo52, and particularly showing a better signal-to-noise ratio compared with the previous GFP-Cdc8 probe. Our investigations also revealed a function for tropomyosin in meiotic actin rings (Yan and Balasubramanian, 2012) in which Cdc8-Tropomyosin has not been previously detected. Because For3 cooperates with the Arp2/3 to assemble the meiotic actin rings (Yan and Balasubramanian, 2012), this suggests that Cdc8 decorates, and perhaps stabilizes the For3-formin-assembled part of the meiotic actin rings.

Fourth, currently, tools composed from native proteins are not available to investigate dynamics of actin cables and the CAR in *S. japonicus*. Despite close phylogeny and morphological similarity to *S. pombe*, CAR assembly in *S. japonicus* is executed through a different, post-anaphase mechanism (Gu et al., 2015). Currently, the proportion of medial CAR actin filaments derived from medial “node-dependent” nucleation versus those transported from non-medial locations is unknown. These can now be investigated.

Finally, although extensive work has been done on *S. cerevisiae* actin cables (Alioto et al., 2016; Chesarone et al., 2009; Huckaba et al., 2004; McInally et al., 2021; Miao et al., 2016; Miao et al., 2013; Yang and Pon, 2002), dynamics of Tpm1 or Tpm2 in interphase cables has not been explored in live cells. Furthermore, an extensive body of work has been done on the interplay between actin patches, actin cables, and endocytosis (Drubin et al., 2005; Kaksonen et al., 2005; Kaksonen et al., 2006). The mNG-Tpm1 and mNG-Tpm2 strains should allow further characterization of such links between actin patches, cables, and endocytosis. The mNG-40L-Tpm1/2 tool can also be used to determine isoform specific localization patterns for Tpm1 and Tpm2 which would help in distinguishing their shared and distinct functions with respect to the S. cerevisiae actin cytoskeleton.

## Materials and Methods

### Construction of *S. pombe* integration plasmid to express mNeonGreen-Cdc8

[Fragment_1] mNeongreen fused with 40 amino acid flexible linker (40 aa linker: LEGSGQGPGSGQGSGSPGSGQGPGSGQGSGPGQGSGPGQG) DNA sequence was PCR amplified from a plasmid template (stored in plasmid collection in Balasubramanian laboratory (pDUAL-Padh21-mNeongreen-40aa, #TH-8-76)) using a pair of PCR primers (forward primer: ATGGTGAGCAAGGGCGAGG; reverse primer: TCCCTGACCGGGGCC).

[Fragment_2] pDUAL vector expressing Cdc8 under the control of the native cdc8 promoter and terminator (used in Palani et al. 2019) were used for the PCR template to PCR amplify the entire sequence of the plasmid using the primers (forward primer: GTTCTGGCCCCGGTCAGGGAATGGATAAGCTTAGAGAGAAAATTAATGCCGC; reverse primer: TCCTCGCCCTTGCTCACCATtttcctactgtttccttctttccttgatgg). Resulting linear DNA fragment contains cdc8 cds at the 5’ and Pcdc8 promoter at the 3’ and both ends were fused with 20 bp overlap sequence with 40 aa linker and mNeongreen, respectively.

[Gibson assembly] (pDUAL:Pcdc8:mNeongreen-40aa:cdc8; #TH8-77) was constructed by Gibson assembly of the Fragment_1 and Fragment_2 above mentioned.

The plasmid #TH8-77 was digested by Not I and DNA fragment containing cdc8 gene fused with mNeongreen and a 40 aa linker were purified by gel extraction (Qiagen). Obtained DNA fragment was used for the transformation of MBY101 strain (*h-, leu1-32, ura4-D18, ade6-210*) and MBY102 strain (*h+, leu1-32, ura4-D18, ade6-210*) by using Lithium acetate method (Alfa C, Fantes P, Hyams J, McLeod M Warbrick E Experiments with fission yeast: a laboratory course manual (Cold Spring Harbor Laboratory Press, Plainview, NY, 1993)). The DNA fragment is integrated at the *leu1-32* gene locus of the cells by endogenous homologues recombination as described previously (Matsuyama et al., Yeast, 2004). Resulting transformant reconstructed *leu1^+^* gene to be leucine prototroph. They were selected on EMM media containing with Uracil/Adenine/Lysine/Histidine and stored as MBY12825 and MBY12828 for *h-* and *h+*, respectively.

### Generation of mNeonGreen tagged *S. japonicus* Tropomyosin

To express a N-terminus fluorescence tagged *S. japonicus* Tropomyosin/Cdc8 (SJAG_04887) as a second copy in *S. japonicus* cells, a DNA fragment of mNeonGreen-40 amino acid linker-cdc8 was synthesised and cloned into pUC-GW-Kan by GeneWiz. This fragment contains the 5’ UTR of *cdc8* between GGGTTTAGTGAG and CAAGAACATCAA, followed by the ORF of mNeonGreen, a 40 amino acid peptide linker (LEGSGQGPGSGQGSGSPGSGQGPGSGQGSGPGQGSGPGQG), the coding sequence of cdc8, and the 3’ UTR of *cdc8* ending at TGTCTTGCTTAG. The DNA fragment of mNeonGreen-40 amino acid linker-cdc8 was subsequently cloned into pSO550 (Gu et al., 2015) flanked by Kpn I and BamH I restriction enzymes. The pSO550-mNeonGreen-40 amino acid linker-cdc8 plasmid was linearised by the restriction enzyme Afe I and integrated at the *ura4sj-D3* locus according to the electroporation protocol described (Aoki et al., 2010).

### Generation of *S. cerevisiae* mNG-Tpm1 and mNG-Tpm2 expressing strains

All primers and plasmids used in this study are listed in Table 1 and Table 2, respectively. Genomic DNA of ESM356 (wild-type) strain was isolated using a LioAc-SDS based protocol as described (Looke et al., 2011). PCR amplification of desired fragments was performed using NEB Q5 High-Fidelity DNA polymerase (#M0491S, NEB). The PCR fragments were assembled into the linearized vector (pRS305; integration vector) (Sikorski and Hieter, 1989) using NEBuilder HiFi DNA Assembly mix (#E2621L, NEB) reaction as per the manufacturer’s instructions. The reaction product was transformed into E. coli TOP10 (#C404010, Invitrogen) cells and plasmid isolation was done using ThermoFisher GeneJet Miniprep Kit (#K0503, ThermoScientific). The positive transformants were confirmed using restriction digestion and sequencing. ESM356 (S288C background) was used as a wild-type strain and all subsequent strains were derived from it. Yeast culturing and transformation was done using established protocols. Yeast expression plasmids carrying mNG-40L-Tpm1/Tpm2 (piSP347 and piSP349 respectively) were linearized with Kas I and transformed for integration into the *leu2* locus in the yeast genome as a second copy. The positive transformants were selected on SC-Leu plates and expression of mNG-40L-Tpm1 and Tpm2 were confirmed using fluorescence microscopy. Yeast strains generated in this study are listed in Table 3.

### Generation of mNeonGreen-Tropomyosin2 expressing RPE-1 cells

mNeonGreen-Tropomyosin2 mammalian expression plasmid construction: a mammalian expression backbone vector for humanized mNeonGreen (mNG), pmNeonGreenHO-G (Addgene, #127912), was linearized using BspE I restriction enzyme at 37°C. A double stranded DNA gblock (IDT) was synthesised consisting of a 40 amino acid flexible linker region (40L: LEGSGQGPGSGQGSGSPGSGQGPGSGQGSGPGQGSGPGQG) fused to the N-terminus of human tropomyosin 2 (TPM2) protein (NCBI, NP_003280.2). Overhang sites complementary to the vector were added to both gblock ends via PCR (forward primer: GGCA TGGACGAACTCTATAAGCTGGAAGGCTCTGGCCAGGGT; reverse primer: ACCGCCTCC ACCGGATCTGAGTTACAAGGATGTTATATCATT) and the fragment was cloned into the vector using NEBuilder^®^HiFi DNA Assembly Master Mix (NEB, #E2621L) to generate a sequence encoding mNG-40L-TPM2 under control of a CMV promoter. Cloning was screened via sequencing (forward primer: CCGGACAATGCAGTTTGAAG) and restriction enzyme digestion.

RPE-1 cell culture: RPE-1 cells were cultured in Dulbecco’s Modified Eagle’s Medium/Nutrient Mixture F-12 Ham with 15 mM HEPES and sodium bicarbonate (Sigma, D6421) supplemented with 6mM L-glutamine (Gibco, 25030-081), 10% foetal bovine serum (Sigma, F7524), 100 U/ml penicillin and 100 μg/ml streptomycin (Gibco, 15140-122) at 37°C under 5% CO_2_.

Transfection: RPE-1 cells were grown on ibiTreat 2 Well μ-Sildes (Ibidi, 80286) to 80% confluency. Cells were then transfected using Lipofectamine^®^2000 (Invitrogen, 11668-019), and Opti-MEM^®^reduced-serum media (Gibco, 31985-062) according to the manufacturer’s instructions.

### Live-cell imaging

#### S. pombe

##### SD confocal microscopy

Andor Revolution XD spinning disk confocal microscopy was used for the image acquisition. The microscope was equipped with a Nikon ECLIPSE Ti inverted microscope, Nikon Plan Apo Lambda 100×/1.45-NA oil-immersion objective lens, a spinning-disk system (CSU-X1; Yokogawa Electric Corporation), and an Andor iXon Ultra EMCCD camera. Images were acquired using the Andor IQ3 software at 69nm/pixel, except in Figure 1D and 2D which were at 80 nm/pixel. The fluorophores were excited by laser lines at wavelengths of 488 or 561 nm. All images were acquired at 25°C.

##### Immunofluorescence imaging

*S. pombe* cells were grown in YEA medium at 24°C to mid-log phase, with OD_595_ 0.2-0.4. 20 mL of cells were collected at 3000 rpm for 2 minutes. Cells were washed with PBS before being fixed in 4% paraformaldehyde at room temperature for 12 minutes. Cells were washed with PBS and resuspended in 200 μL protoplasting solution. Protoplasting solution was made with 1mg/mL lysing enzyme (Sigma, L1412) and 6 μL/mL Zymolyase (GBiosciences, 1.5 U/μL) in PBS with 1.2 M sorbitol. The cells were incubated at room temperature for 15 minutes and visually checked for protoplasting by mixing an aliquot 1:1 with 10% SDS. To inactivate the protoplasting enzymes, 1mL of 1% Triton was added and incubated for 2 minutes. Cells were then spun down and blocked by resuspension in 0.5 mL PBAL (10% BSA, 10 mM lysine hydrochloride, 50 ng/mL carbenicillin and 1 mM sodium azide) and incubated at room temperature for 1 hour, gently rocking. After centrifugation, primary, affinity-purified anti-cdc8p was added (1:200 in PBAL) and incubated at 4°C overnight. Cells were washed twice in PBAL. Secondary antibody (Life Technologies, 1420898) was added (1:200 in PBAL) for 90 minutes at room temperature. Following two washes the cells were ready for imaging.

###### Sample preparation for live-cell imaging

*S. pombe* cells were grown in YEA medium at 24°C to mid-log phase. 20 mL of cells were collected at 3000 rpm for 1 minute and added on 2% agarose in YEA medium pad and sealed with VALAP.

For Figure 2A, 2B, 2C and 2E, *S. pombe* cells were grown in YEA medium at 24°C to mid-log phase. 20 mL of cells were spun down and added on 2% agarose in YEA medium pad before sealing with VALAP. Time-lapse images were acquired at 3 seconds interval. A total of 16 planes were imaged with Z-step sizes of 0.4 μm.

For Figure 4C, time-lapse images were acquired at 1 minute interval. A total of 13 planes were imaged with Z-step sizes of 0.5 μm.

For Supplemental figure 3, *S. pombe cdc25-22* cells were grown in YEA medium at 24°C to midlog phase. The cells were blocked at the G2/M transition by incubating at 36°C for 4 hours. 20mL of cells were collected and added on 2% agarose in YEA medium pad before sealing with VALAP. Time-lapse imaging commenced about 15 minutes after cells were removed from 36°C into 25°C for synchronous release into the cell cycle. Time-lapse images were acquired at 2 minutes interval. A total of 17 planes were imaged with Z-step sizes of 0.5 μm.

###### For ring isolation and ATP addition

*S. pombe* cells were grown in YEA medium at 24°C to mid-log phase, with OD_595_ 0.2-0.4. 20 mL of cells were collected at 3000rpm for 1 minute. Cells were washed with equal volume of 1X E-Bufffer (50 mM sodium citrate and 100 mM sodium phosphate, pH6.0) before resuspending in 5 mL of 1X E-buffer supplemented with 1.2 M sorbitol. 0.6 mg/mL of lysing enzyme (Sigma, L1412) was added. Cells were incubated at 24°C, shaking at 80 rpm. After 1.5 hour, 30 μL of LongLife Zymolyase (GBiosciences, 1.5U/μL) was added, and cells were incubated at 24°C for another 1.5 hour. The following washes were done with centrifugation at 450xg for 2 minutes. Protoplasts were washed with 1X E-Buffer with 0.6 M sorbitol then resuspended in EMMA supplemented with 0.8 M sorbitol. The protoplasts were incubated at 24°C, shaking at 80 rpm for 3.5 hours. The following steps were done on ice. After a wash with wash buffer, cell ghosts were obtained by permeabilizing the protoplasts with isolation buffer containing 0.5% NP-40. The cell ghosts were then homogenized in a glass homogenizer to obtain isolated rings. The rings were washed and resuspended in reactivation buffer. Imaging of isolated rings was done using a CellASIC ONIX Microfluidic system (Merck Millipore) with 0.5 mM ATP in reactivation buffer.

##### Spot assay

*S. pombe* cells were cultured in YEA media at 24°C to saturation, subjected to six-fold serial dilution, and spotted onto YEA agar. Plates were incubated at 24°C and 36°C for 3 days before being photographed.

##### Image analysis

Images were viewed and analysed using ImageJ. All image stacks were projected along the Z axis (maximum intensity) for analysis and for representation. If applicable, the movies were bleach corrected using “Image/Adjust/Bleach Correction” in ImageJ. The background of all microscopy images was subtracted in “Image/Adjust/Brightness & contrast”. All time-lapse videos were edited by ImageJ and saved in MP4 format with H.264 compression.

##### *Schizosaccharomyces pombe* mating and sporulation

Homothallic (h90) strains able to switch mating types were used, where cells were grown in liquid or agar Minimum Sporulation Media (MSL), with or without nitrogen (+/-N) (Egel et al., 1994). Live imaging of *S. pombe* mating cells protocol was adapted from (Vjestica et al., 2016). Briefly, cells were first pre-cultured overnight in MSL+N at 25°C, then diluted to OD_600_ = 0.05 into MSL+N at 25°C for 20 hours. Exponentially growing cells were then pelleted, washed in MSL-N by 3 rounds of centrifugation, and resuspended in MSL-N to an OD_600_ of 1.5. Cells were then grown 3 hours at 30°C to allow mating in liquid, added on 2% agarose MSL-N pads, and sealed with VALAP. We allowed the pads to rest for a minimum of 30 min at 30°C before imaging.

Images presented in Figures 1E and S2 were obtained using a ZEISS LSM 980 scanning confocal microscope with 4 confocal Detectors (2x GaAsP, 2x PMT), an Airyscan2 detector optimized for a 60x/1.518 NA oil objective, and 6 Laser Lines (405nm, 445nm, 488nm, 514nm, 561nm, 640nm) on inverted Microscope Axio Observer 7. Images were acquired using the Airyscan2 detector and processed with the Zen3.3 (blue edition) software for super resolution.

Images presented in Figure 2F were obtained using a DeltaVision platform (Applied Precision) composed of a customized inverted microscope (IX-71; Olympus), a UPlan Apochromat 100×/1.4 NA oil objective, a camera (CoolSNAP HQ2; Photometrics or 4.2Mpx PrimeBSI sCMOS camera; Photometrics), and a color combined unit illuminator (Insight SSI 7; Social Science Insights). Images were acquired using softWoRx v4.1.2 software (Applied Precision). Images were acquired every 5 minutes for 12 hours. To limit photobleaching, overnight videos were captured by optical axis integration (OAI) imaging of a 4.6 μm z-section, which is essentially a real-time z-sweep.

Images presented in Figure 5D were obtained using a spinning-disk microscope composed of an inverted microscope (DMI4000B; Leica) equipped with an HCX Plan Apochromat 100×/1.46 NA oil objective and an UltraVIEW system (PerkinElmer; including a real-time confocal scanning head [CSU22; Yokagawa Electric Corporation], solid-state laser lines, and an electronmultiplying charge coupled device camera [C9100; Hamamatsu Photonics]). Images were acquired using the Volocity software (PerkinElmer).

#### Schizosaccharomyces japonicus

Cells expressing mNeonGreen-cdc8 were grown in YES medium to OD_595_ 0.4-0.6 at 30 °C. Prior to imaging, 1 mL yeast culture was concentrated to 50 μL after centrifugation at 1500x g, 30 sec. 1 μL cell suspension was loaded on YES agarose pad (2% agarose) and covered with 22 x 22 mm glass coverslip (VWR #631-0125, thickness: 1.5). Time-lapse imaging of mNG-cdc8 cells were performed at 30 °C with acquisition of 16 z-slices, 0.6 μm per slice and 11.2 seconds per time interval.

Temperature-sensitive *cdc25-D9* cells expressing mNeonGreen-cdc8 were grown in YES medium to OD_595_ 0.1 at 24 °C. 50 mL cell culture was incubated at 36 °C for 3 hours 15 minutes to block cell cycle progression of these cells at G2/M transition before cells were released to permissive temperature, allowing synchronised mitotic entry of *cdc25-D9* cells. Time-lapse imaging of mNG-cdc8 expressing *cdc25-D9* cells were performed at 25 °C with acquisition of 13 z-slices, 0.6 μm per slice and 30 seconds per time interval.

*S. japonicus* cells expressing mNG-cdc8 were grown in EMM liquid media adequately complemented at 30°C to exponential phase, then washed in SPA liquid medium by 3 rounds of centrifugation, added on 2% agarose SPA pads, and sealed with VALAP. Cells were then allowed to mate for 7 hours before imaging.

Spinning-disk confocal images are acquired with iXon Ultra U3-888-BV monochrome EMCCD camera (Andor Technology Ltd., UK), Eclipse Ti-E inverted microscope (Nikon, Japan) fitted with CSU-X1 spinning disk confocal scanning unit (Yokogawa electric, Japan), 600 series SS 488 nm, 50 mW laser, Brightline single band filter FF01-525/50-25 (Semrock, USA), CFI Plan Apo Lambda 100× (N.A. = 1.45) oil objective (Nikon, Japan). Image acquisition and deconvolution are controlled by Andor Fusion software (2.3.0.36).

#### Saccharomyces cerevisiae

Glass bottom dishes (35mm; #81218, Ibidi GmBH, Germany) were coated with 6% concanavalin A (#C2010, Sigma). Yeast cells grown till mid-log phase in filter-sterilized SC-complete medium at 30°C were plated on the coated dish for imaging. Time-lapse Imaging was performed at 30°C using Andor Revolution XD spinning disk confocal microscopy. The microscope was equipped with a Nikon ECLIPSE Ti inverted microscope, Nikon Plan Apo Lambda 100×/1.45-NA oil-immersion objective lens, a spinning-disk system (CSU-X1; Yokogawa Electric Corporation), and an Andor iXon Ultra EMCCD camera. The fluorophores were excited by laser lines at wavelengths of 488 nm. Figure S6C images were acquired using Andor Dragonfly 502 spinning disk confocal system, (Andor Technology Ltd., Belfast, UK) equipped with Andor Sona scMOS camera mounted on a Leica Dmi8 inverted fully motorized microscope (Leica, Wetzlar, Germany). Images acquired comprised of 18 z-slices spaced 0.35μm, using a 100x oil objective (1.4 NA) and solid-state lasers 488nm (mNG) for excitation. Images were deconvolved using Andor Fusion software (2.3.0.44) and processed offline using Fiji (Schindelin et al., 2012).

### Human RPE cells

SD confocal microscopy: Images were acquired via the same microscopy system as the *S. pombe* images (see above), with a Nikon Plan Fluor 40×/1.30 oil immersion objective lens additionally used. Images were acquired at 80 nm/pixel and fluorophores were excited by 488 or 561 nm lasers.

Sample preparation for live-cell imaging: Transfected RPE-1 cells were imaged in phenol-free Leibovitz’s L-15 Medium (Gibco, 21083-027) at 37°C.

Sample preparation for Rhodamine phalloidin co-stained fixed-cell imaging: Transfected RPE-1 cells were fixed in 4% paraformaldehyde/PBS, stained with Rhodamine phalloidin (Invitrogen, R415) diluted 1:400 in 0.1% Triton X-100/PBS and sealed with Vectashield^®^(Vector, H-1000). Images were acquired at room temperature.

### Tropomyosin nanobody construction

Tropomyosin (SpCdc8) was cloned into pET^MCN^ vector (without any tag) for protein expression as described previously (Kompula et al., 2012; Palani et al., 2019). Purified recombinant SpCdc8 was dialyzed against the tropomyosin storage buffer (50 mM NaCl, 10 mM imidazole, pH 7.5, and 1 mM DTT) and flash frozen in liquid nitrogen. It was sent to Hybrigenics to raise nanobodies against *S. pombe* Cdc8. The nanobody protein sequences provided by Hybrigenics were codon optimized for *S. pombe* expression and gBlocks were synthesized (IDT, US). The nanobody gBlocks were cloned into *S. pombe* vector (pDUAL) (Matsuyama et al., 2008) under the constitutive expression promoter (pADH11) in fusion with mNeonGreen (referred to as mNG) along with 40 aa Linker at the N-terminus of nanobodies. Yeast expression nanobody plasmids were linearized with Not I restriction enzyme and transformed into MBY 192 for integration. Colonies were selected on the EMM-Leu plates and nanobody expression was confirmed by fluorescence microscopy.

### Yeast Two-Hybrid

Indicated genes of interest: Tropomyosin (SpCdc8) from *S pombe* genomic DNA and S. pombe codon optimized nanobodies (A5, A10, A19, A22, A37, A83 and A94) from IDT gblocks were amplified and cloned into pMM5S (activation domain of Gal4p) and pMM6S (DNA binding protein LexA) plasmids, respectively. Plasmids carrying the genes of interest (SpCdc8, 7 nanobodies) and empty vector controls were transformed into the yeast strains SGY37 (MATa) and YPH500 (MATα). Transformants with the desired plasmids were selected on SC plates lacking Histidine (-His) or Leucine (-Leu), respectively. Mating of the MATa and MATα was done on YPD plates which were incubated for two days at 30 □C, followed by replica-plating on double selection SC-His-Leu plates. The plates were incubated for two days at 30 □C before the X-Gal (#RC-212, G-Biosciences, USA) overlay. X-Gal overlay assay was performed as described (Geissler et al., 1996). Plates were scanned after 24 hr of incubation with the X-Gal overlay mixture.

### AlphaFold 2 prediction

For prediction of the Cdc8-nanobody complex, two Cdc8 (CAA93291.2) and one Nb5 sequence or two Cdc8 and one BC2-nanobody (PDB:5IVO) (Braun et al., 2016) sequences were input on AlphaFold Colab (https://colab.research.google.com/github/deepmind/alphafold/blob/main/notebooks/AlphaFold.ipynb). The predicted structures were visualized and overlaid as to Cdc8 in UCSF Chimera version 1.14 (https://www.rbvi.ucsf.edu/chimera) (Pettersen et al., 2004).

## Supporting information

Yeast Strains

## Acknowledgements

Funding: Mohan Balasubramanian: Wellcome Collaborative (203276/Z/16/Z), Wellcome Senior Investigator (WT101885MA), European Research Council (AdG-Actomyosin Ring)

Funding: Sophie Martin: Swiss National Science Foundation (310030B_176396) and European Research Council (CoG CellFusion)

Funding: Snezhana Oliferenko Wellcome Trust Investigator Award in Science (220790/Z/20/Z) and BBSRC (BB/T000481/1).

Funding: Saravanan Palani: DBT-Wellcome Trust India Alliance Intermediate fellowship grant (IA/I/21/1/505633) and DST-SERB-(SRG/2021/001600) and GATE fellowship (for AD).

## Supplemental Figure Legends

**Supplementary figure 1: Tagging Cdc8 with mNeonGreen in *S. pombe***. (A) Schematic illustration showing the insertion of P*_cdc8_* mNG-40 amino acid linker-cdc8 cassette at *leu1* locus. (B) Growth of strains was assessed by manual spot test. Untagged wild-type strains act as a control for mNG-Cdc8 strains. Cells were cultured in YEA media at 24°C to saturation, subjected to six-fold serial dilution, and spotted onto YEA agar. Plates were incubated at 24°C and 36°C for 3 days before being photographed. (C) Fimbrin null mutant cells (*fim1*Δ) cells show obvious mNG-Cdc8 patches.

**Supplementary figure 2: The mNG-Cdc8 probe is more sensitive than the previously used GFP-Cdc8 probe**.Airyscan2 images of mating cells expressing GFP-Cdc8 (top; green) or mNG-Cdc8 (bottom; green) and Myo52-tdTomato (magenta), which labels the fusion focus. mNG-Cdc8 allows detection of actin cables not visible with GFP-Cdc8. Higher cytosolic GFP signals were detected in the GFP-cdc8 cells. Scale bars are 5μm.

**Supplementary figure 3: Actomyosin ring assembly visualized using mNG-Cdc8 in elongated *cdc25-22* cells**.Panel i and ii are time-lapse images of two *S. pombe cdc25-22* cells expressing mNG-Cdc8 demonstrating medial assembly of Cdc8 cables as well as flow of non-medial cables containing Cdc8-tropomyosin into the CAR. Scale bar is 5μm.

**Supplementary figure 4: Tagging Cdc8 with mNG in *S. japonicus***. (A) Schematic illustration showing the insertion of P*_cdc8_* mNG-40 amino acid linker-*cdc8* cassette at the *ura4* locus. The tag was expressed under the native cdc8 promoter. (B) Growth of strains was assessed by manual spot test. Untagged wild-type strains act as a control for mNG-Cdc8 expressing strains. Cells were cultured in YEA media at 24°C to saturation, subjected to six-fold serial dilution, and spotted onto YEA agar. Plates were incubated at 24°C and 36°C for 3 days before being photographed.

**Supplementary figure 5: Actomyosin ring assembly visualized using mNG-Cdc8 in elongated *S. japonicus cdc25-D9* cells**.Time-lapse images of an *S. japonicus cdc25-D9* cell expressing mNG-Cdc8 demonstrating medial assembly of Cdc8 cables as well as flow of non-medial cables containing Cdc8-tropomyosin into the CAR. Scale bar is 5μm.

**Supplementary figure 6: Tagging Tpm1 and Tpm2 with mNG in *S. cerevisiae***. (A) Schematic illustration showing the insertion of P*_Tpm1/2_* mNG-40 amino acid *linker-Tpm1/2* cassette at the *leu2* locus. (B) Growth of strains was assessed by manual spot test. Untagged wild-type strains act as a control for mNG-tagged Tpm1 and Tpm2 strains. Cells were cultured in YPD media at 24°C to saturation, subjected to six-fold serial dilution, and spotted onto YPD agar. Plates were incubated at 23°C and 37°C for 3 days before being photographed. (C) Fimbrin null mutant cells (*sac6*Δ cells) show obvious mNG-Tpm1 and mNG-Tpm2 patches. Scale bar is 5μm.

**Supplementary figure 7: mNG-TPM2 in human RPE-1 cells**. (A) Schematic illustration showing the expression of mNG-40 aa linker-TPM2 under the control of a CMV promoter in an RPE-1 cell. (B) Fixed cells expressing mNG-TPM2 show mNG colocalization with Rhodamine phalloidin-stained F-actin. Scale bar is 5μm.

**Supplementary figure 8: Nanobody-mNeonGreen in *S. pombe***. (A) Yeast two hybrid interaction shown between Sp Cdc8 and 7 camelid nanobodies. (B) Schematic illustration showing the insertion of Nb5-40 aa linker-mNG cassette at the *leu1* locus. (C) Growth of strains was assessed by manual spot test. Untagged wild-type strains act as a control for Nb5-mNG strains. Cells were cultured in YEA media at 24°C to saturation, subjected to six-fold serial dilution, and spotted onto YEA agar. Plates were incubated at 24°C and 36°C for 3 days before being photographed.

## Movie Legends

Movie 1: *S. pombe* mNG-Cdc8 patch and cable dynamics.

Movie 2: mNG-Cdc8 dynamics during CAR assembly in *S. pombe*

Movies 3 and 4: mNG-Cdc8 dynamics during CAR assembly in highly elongated *S. pombe cdc25-*22 cells, demonstrating flow of Cdc8 cables into the CAR during its assembly.

Movie 5: mNG-Cdc8 dynamics and cable expulsion during CAR constriction in *S. pombe*.

Movie 6: Dynamics of mNG-Cdc8 during ATP-dependent constriction of CARs within *S. pombe* cell ghosts.

Movie 7: Dynamics of *S. pombe* mNG-Cdc8 during mating and sporulation.

Movie 8: Dynamics of *S. japonicus* mNG-Cdc8 during CAR assembly and constriction.

Movie 9: Dynamics of *S. japonicus* mNG-Cdc8 during CAR assembly and constriction in elongated *cdc25*-D9 cells.

Movie 10: Dynamics of *S. cerevisiae* mNG-Tpm1 in cables.

Movie 11: Dynamics of *S. cerevisiae* mNG-Tpm1 in the CAR.

Movie 12: Dynamics of *S. cerevisiae* mNG-Tpm1 in cables.

Movie 13: Dynamics of *S. cerevisiae* mNG-Tpm1 in the CAR.

Movie 14: Simulation of Cdc8-Nanobody 5 interaction with Cdc8 dimer.

Movie 15: Dynamics of mNG-Nb5 during CAR assembly in *S. pombe*.

