## Supplementary material for "mNeonGreen-tagged fusion proteins and nanobodies reveal localization of tropomyosin to patches, cables, and contractile actomyosin rings in live yeast cells": Yeast Strains

|  | **Strain number** | **Genotype** |
| --- | --- | --- |
| *S. pombe* | MBY101 | *ade6-210 ura4-D18 leu1-32* h- |
|  | MBY102 | *ade6-210 ura4-D18 leu1-32* h+ |
|  | MBY192 | *ura4-D18 leu1-32* h- |
|  | MBY12825 | *leu1-32<pDUAL:pcdc8:*mNeongreen-40aa:*cdc8* *ura4-D18* *ade6-210, cdc8+* h- |
|  | MBY12828 | *leu1-32<pDUAL:pcdc8:*mNeongreen-40aa:*cdc8 ura4-D18 ade6-210, cdc8+* h+ |
|  | MBY12947 | *leu1-32<pDUAL-pcdc8:*mNeongreen-40aa:*cdc8 lys1<<pLYS1U-Pact1:*lifeact-mCherry:ura4+ *ade6-210*, cdc8+ |
|  | MBY12994 | *fim1∆ leu1-32<pDUAL:pcdc8*:mNeongreen-40aa:*cdc8, ura4-D18, ade6-210,* cdc8+ |
|  | MBY13071 | *cdc25*-22 *leu1-32<pDUAL:pcdc8*:mNeongreen-40aa:*cdc8* *rlc1-*mCherry:*ura4+* *ade6-210* h+ |
|  | MBY13185 | *leu1-32<padh11*:Nanobody5-mNeonGreen |
|  | YSM3935 | h90 myo52-tdTomato:natMX leu1-32:p^cdc8^:mNeonGreen-cdc8:termcdc8:term^ADH1^:leu1+ lys3+:p^map3^:mCherry:term^ADH1^:bsdMX ade6-M216 ura4-D18 |
|  | YSM3936 | h90 myo52-tdTomato:natMX leu1-32:p^ADH1^:cdc8Nb5-mNeonGreen:term^ADH1^:leu1+ ade6-M210 ura4-294 |
|  | YSM3316 | h90 myo52-tdTomato:natMX leu1-32:p^ADH1^:cdc8Nb5-mNeonGreen:term^ADH1^:leu1+ ade6-M210 ura4-294 |
| *S. japonicus* | SOJ5 | matsj-P2028 h- |
|  | SOJ4909 | *pcdc8*-mNeonGreen-40 a.a. linker-cdc8^ORF^-*cdc8^3'UTR^*::ura4+::ura4sj-D3 h+ |
|  | SOJ5001 | *pcdc8*-mNeonGreen-40 a.a. linker-cdc8^ORF^-*cdc8^3'UTR^*::ura4+::ura4sj-D3 cdc25-D9:kanR:ura4+ |
|  | SOJ5221 | *pcdc8*-mNeonGreen-40 a.a. linker-cdc8^ORF^-*cdc8^3'UTR^*::ura4+::ura4sj-D3 h- |
| *S. cerevisiae* | YSP002 | ESM356 MATa ura3-52 leu2∆1 trp1∆63 his3∆200 (wild type) |
|  | YSP107 | As YSP002 except pRS305-Ptpm1-mNG-40aaL-tpm1-Ttpm1-leu2 |
|  | YSP108 | As YSP002 except pRS305-Ptpm2-mNG-40aaL-tpm2-Ttpm2-leu2 |
|  | YSP191 | As YSP002 except pRS305-Ptpm1-mNG-40aaL-tpm1-Ttpm1-leu2 Δsac6::His3MX6 abp1-tdtomato::natNT2 |
|  | YSP192 | As YSP002 except pRS305-Ptpm2-mNG-40aaL-tpm2-Ttpm2-leu2 Δsac6::His3MX6 abp1-tdtomato::natNT2 |
