## Supplementary figures and images for "mNeonGreen-tagged fusion proteins and nanobodies reveal localization of tropomyosin to patches, cables, and contractile actomyosin rings in live yeast cells"

### Supplemental Figure 1

**Supplemental figure 1**

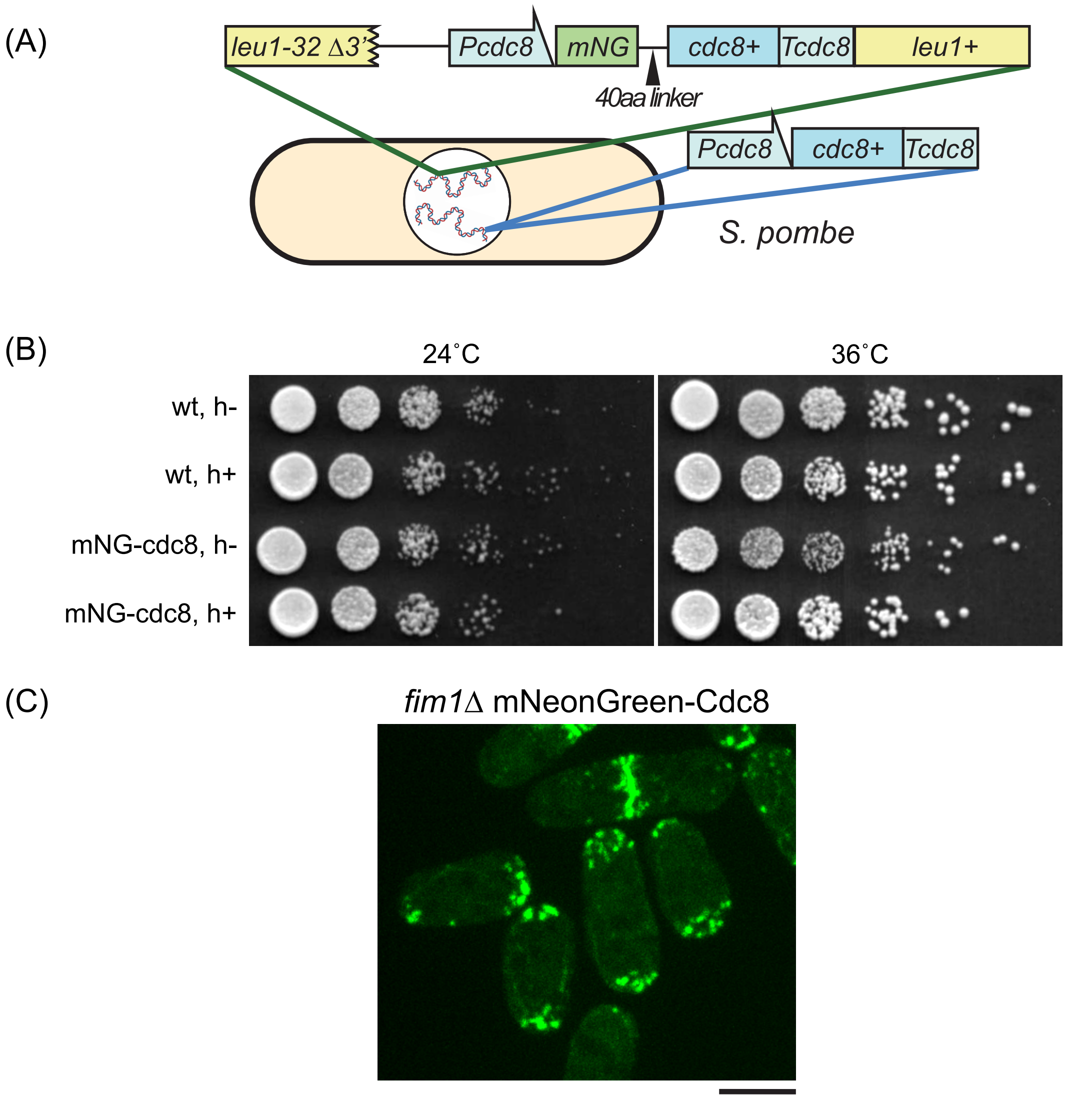

### Supplemental Figure 2

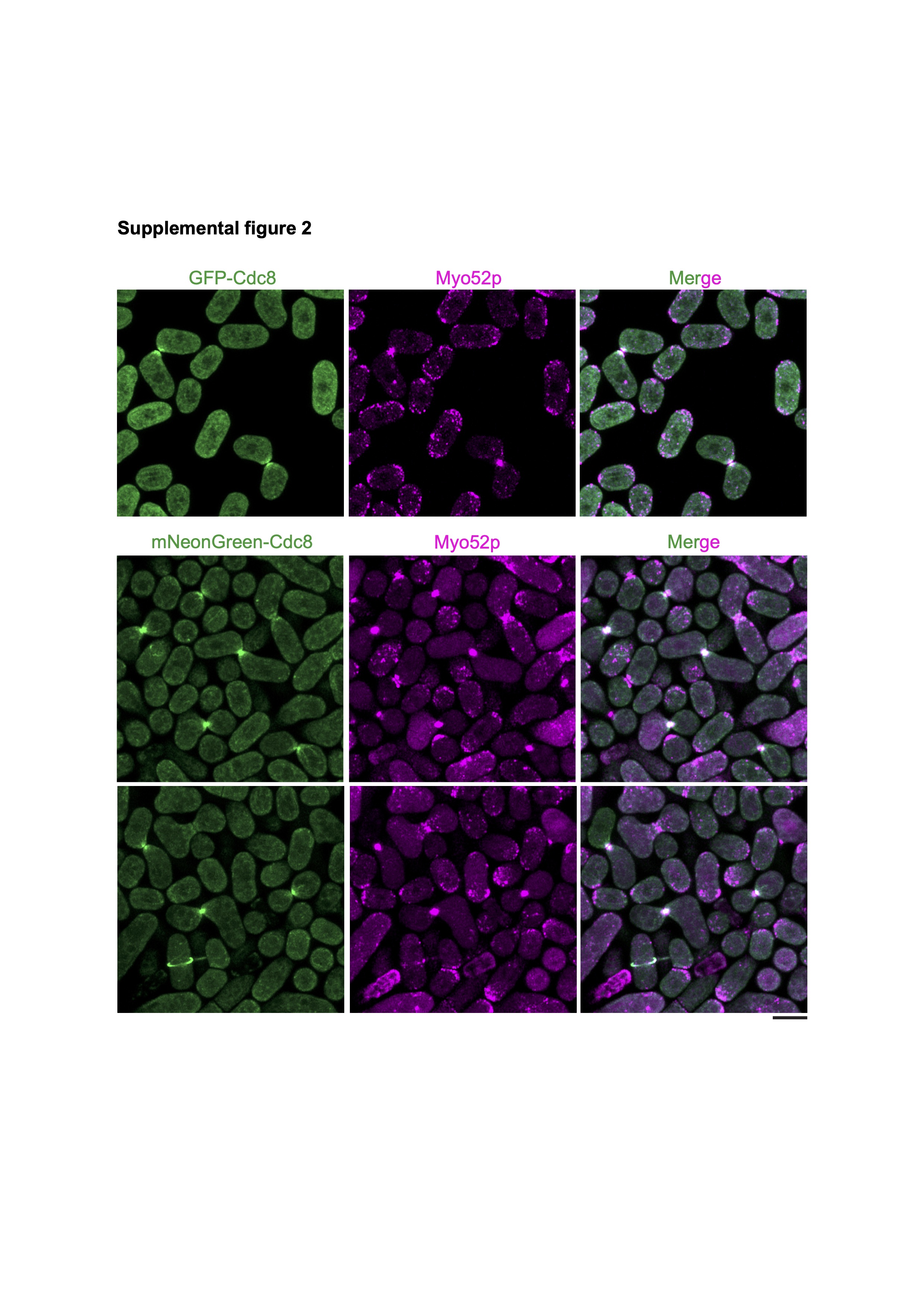

### Supplemental Figure 3

**Supplemental figure 3**

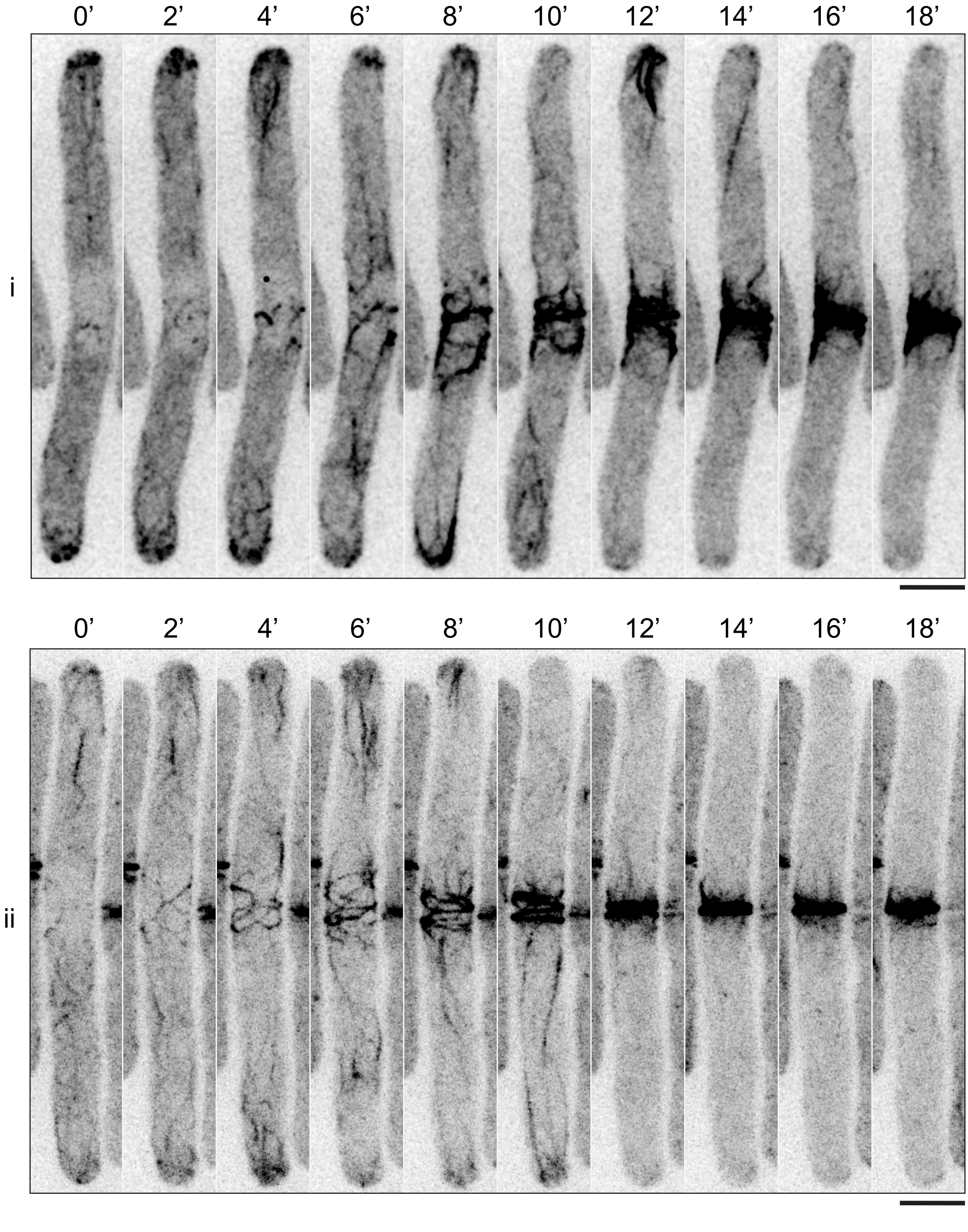

### Supplemental Figure 4

#### Supplemental figure 4

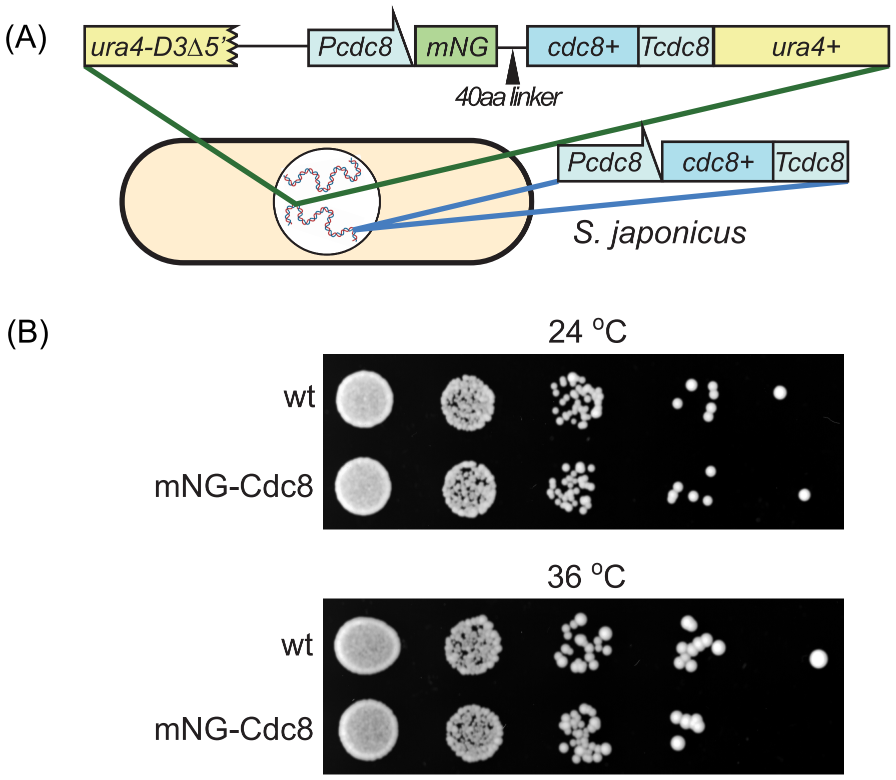

### Supplemental Figure 5

#### Supplemental figure 5

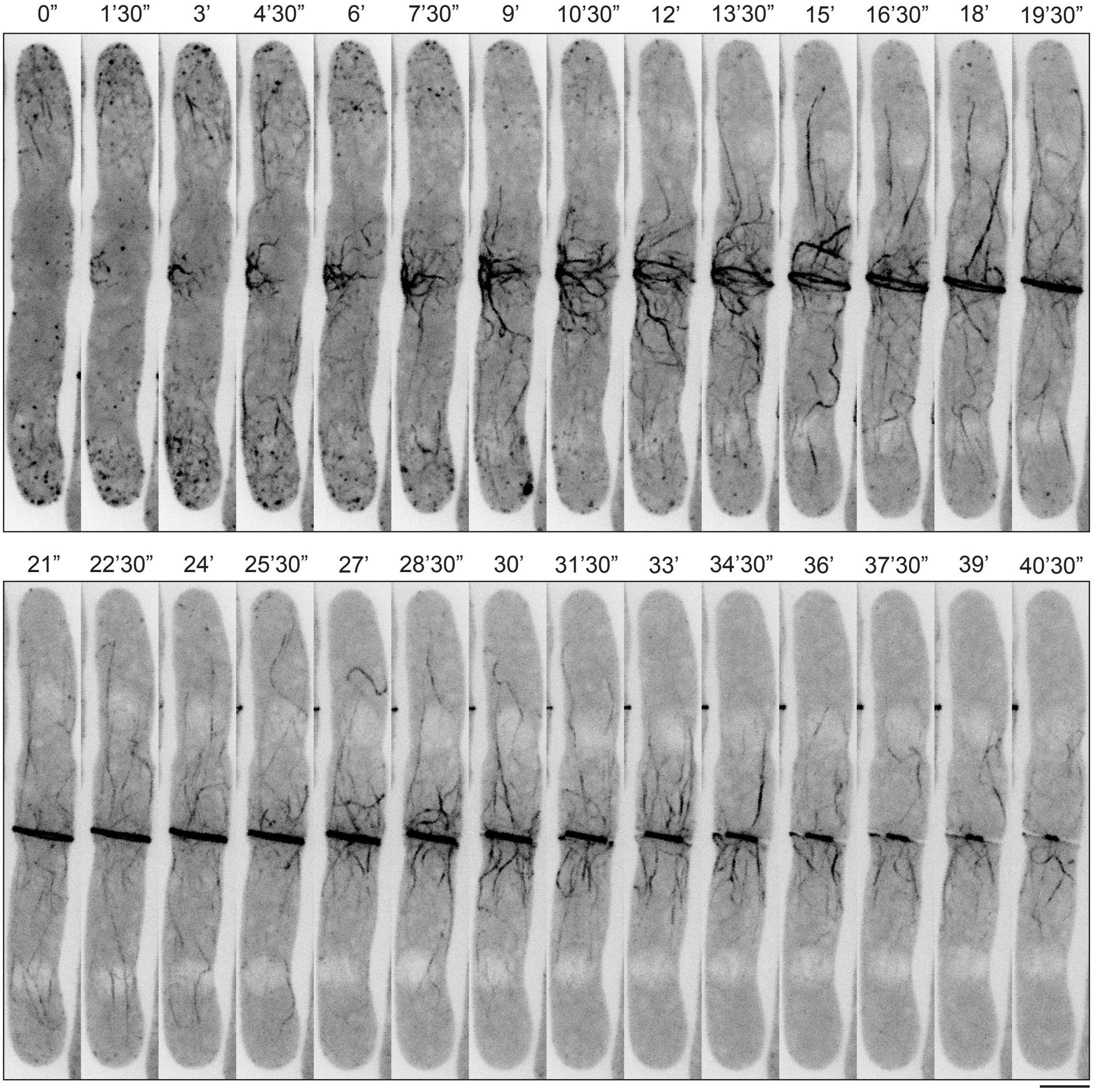

### Supplemental Figure 6

**Supplemental figure 6**

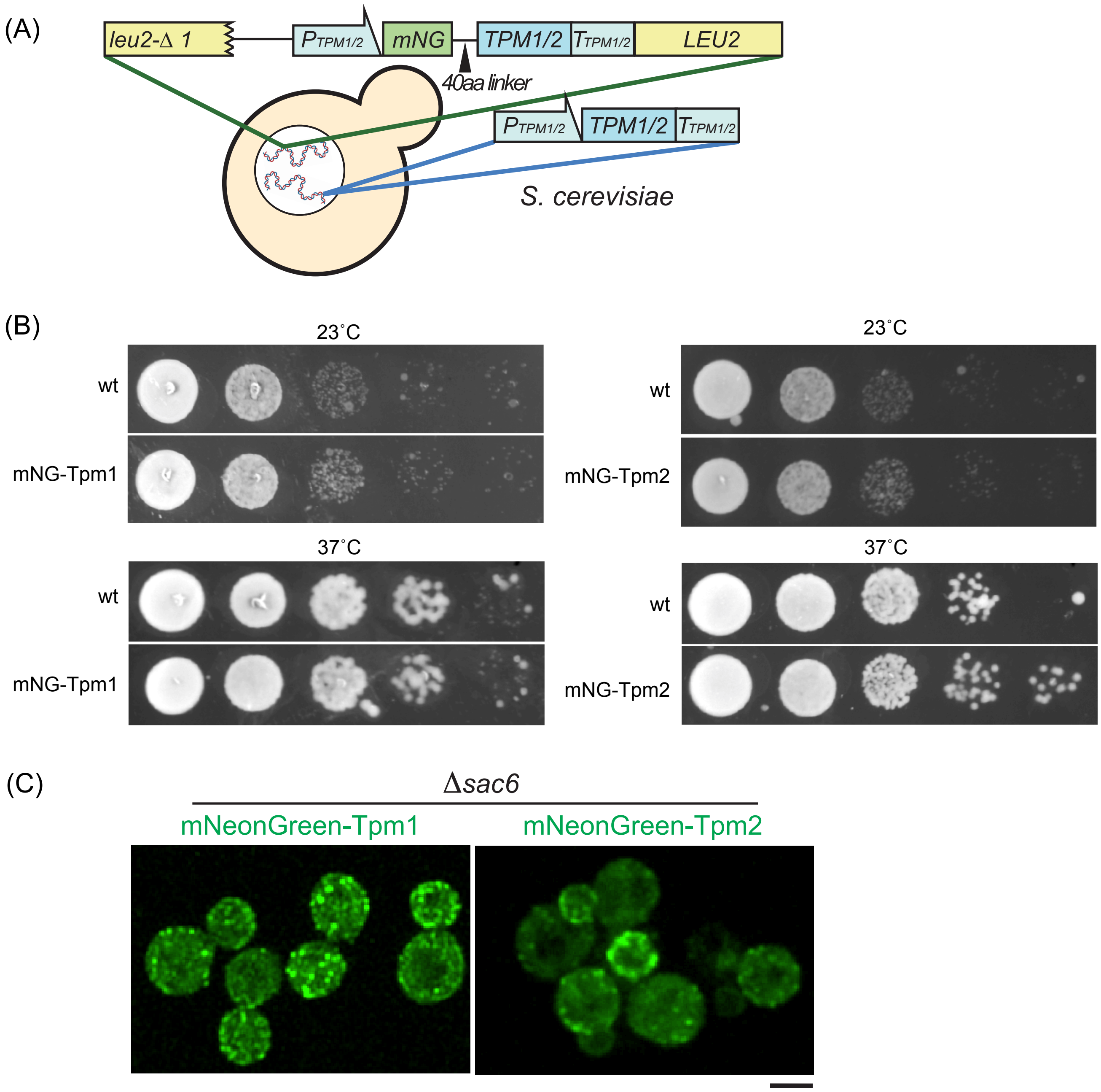

### Supplemental Figure 7

**Supplemental figure 7**

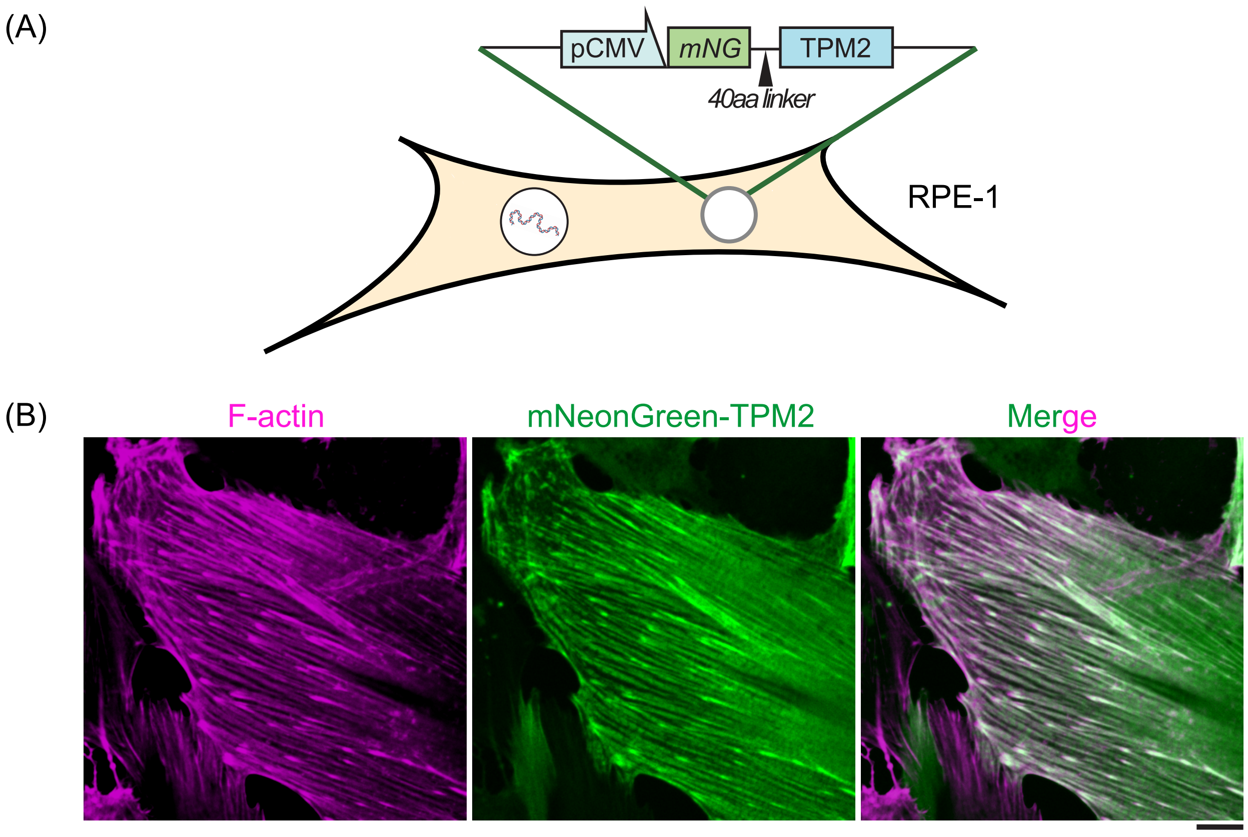

### Supplemental Figure 8

#### Supplemental figure 8

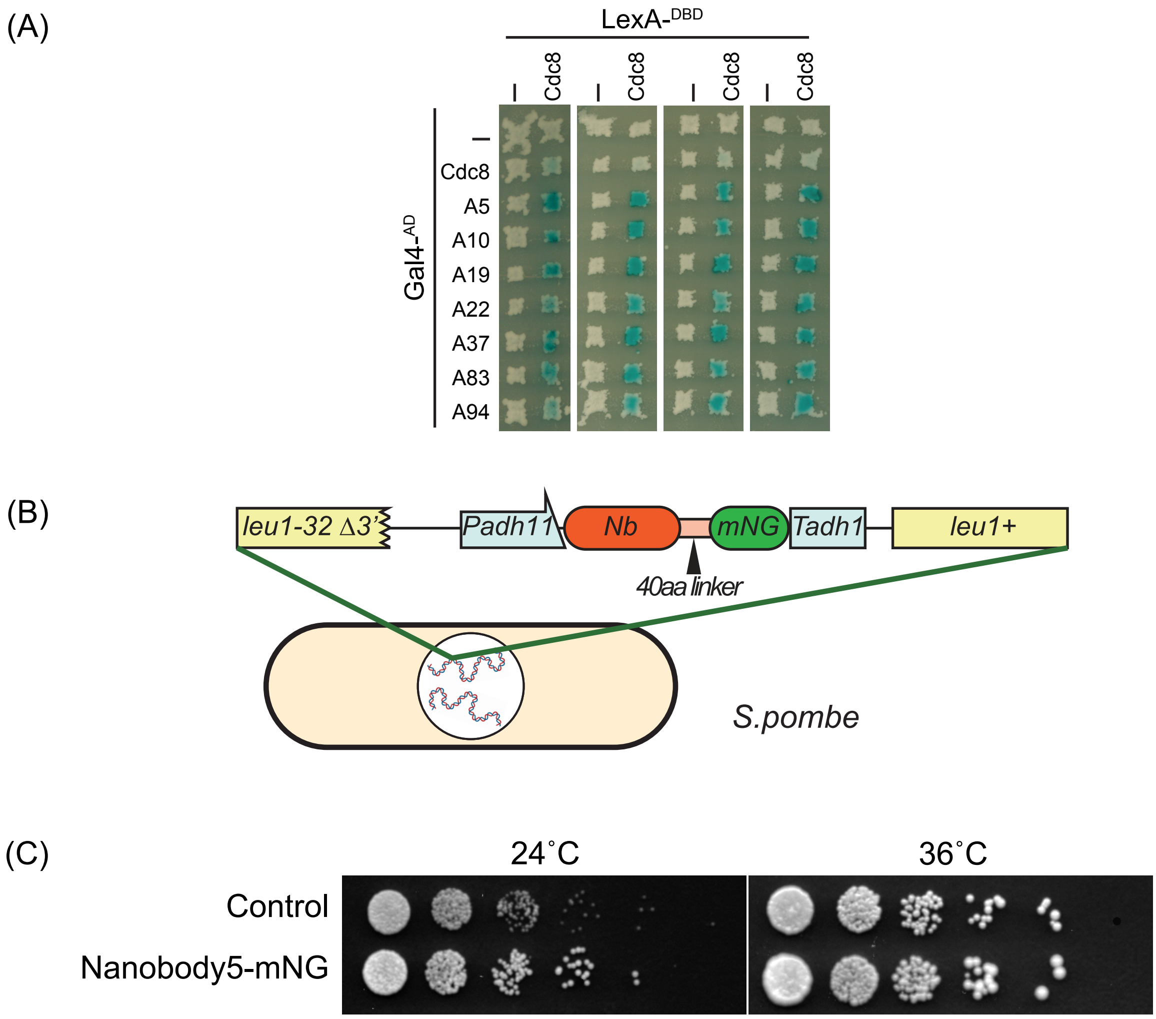
